# Visualizing single-base RNA mutations in living cells through DNA nanostructure-mediated amplification

**DOI:** 10.64898/2026.02.14.705875

**Authors:** Xiao-Yi Ma, Musitaba Mutailifu, Ying Lin, Jian-Heng Qiu, Jun-Jie Wang, Zheng Wu, Yu-Zheng Gan, Lei Zhu, Li-Peng Hu, Qing Li, Jia-Mei Luo, Dong-Xue Li, Zhi-Gang Zhang

**Author notes:** Corresponding authors: Zhi-Gang Zhang,; Dong-Xue Li,; Jia-Mei Luo,. Xiao-Yi Ma, Musitaba Mutailifu, Ying Lin contributed equally to this work.

## Abstract

Capturing RNA dynamics in living cells would provide critical insights into transcriptional control and cellular adaptation, but remains technically formidable — particularly at single-base precision. Here, we introduce a DNA tetrahedron based three-dimensional catalytic hairpin assembly (3D@CHA) nanoplatform that couples target recognition with catalytic activation in a spatially organized framework. Three cascaded hairpins (H-AN, H1, and H2) then enable localized and efficient signal amplification. Without external carriers or transfection, the platform exhibits robust biocompatibility, distinguishing highly homologous *insulin I* (*Ins1*) and *insulin II* (*Ins2*) mRNAs in living cells and tracking their redistribution and intercellular transfer during metabolic changes. Introducing a single-base mismatch site into H1 and coupling it with a Förster resonance energy transfer (FRET) readout yielded a *KRAS*-3D@CHA probe capable of detecting *KRAS^G12D^* mutations at the RNA level with single-base resolution. This platform establishes a programmable framework for precise RNA imaging and mutation discrimination, opening new avenues for RNA-level diagnostics and precision oncology.

## Introduction

RNA orchestrates gene expression, regulates cellular physiology, and underpins diverse biological processes^1–3^. Among its many forms—messenger RNA (mRNA), microRNA (miRNA), long noncoding RNA (lncRNA), circular RNA (circRNA), and transfer RNA (tRNA)—each plays a critical role in maintaining cellular homeostasis and regulating gene networks^4–8^. Abnormal RNA expression, modification, or localization is closely associated with the onset and progression of cancer, neurodegenerative disorders, and metabolic diseases^9–11^. Therefore, real-time visualization and quantification of RNA molecules in living cells are essential for decoding transcriptional dynamics, mapping subcellular localization, and elucidating RNA-mediated regulation in health and disease. This underscores the urgent need for RNA detection technologies that are highly sensitive, specific, real-time, and noninvasive.

Conventional methods such as quantitative PCR (qPCR) and fluorescence *in situ* hybridization (FISH) offer high sensitivity and specificity but require cell lysis or fixation, precluding live-cell monitoring^12,13^. More recent approaches—including MS2/PP7 tagging, CRISPR-Cas-based imaging, and fluorescent RNA aptamers—enable live-cell studies but still face key limitations: they require modification of target RNAs, which may disrupt native function; probes often suffer from poor stability and low signal-to-noise ratio; and some systems can be cytotoxic^14–16^. Thus, existing tools cannot simultaneously achieve sensitivity, specificity, and biological fidelity, particularly for distinguishing homologous transcripts or single-nucleotide variants.

Structured DNA nanotechnology offers a powerful route to overcome these limitations. DNA tetrahedra, assembled from four short oligonucleotides, form rigid three-dimensional scaffolds whose double-stranded edges confer high mechanical stability and nuclease resistance^17^. Their nanoscale geometry enables efficient cellular uptake without transfection agents, while programmable vertices allow precise functionalization with fluorophores, aptamers, or recognition motifs^18–20^. These properties render DNA tetrahedra an ideal structural chassis for constructing robust and modular intracellular sensing platforms.

Catalytic hairpin assembly (CHA) is an enzyme-free isothermal amplification strategy based on toehold-mediated strand displacement^21^. In CHA, single-stranded oligonucleotides fold into stem-loop structures to form hairpin probes. When a target RNA hybridizes with the toehold region of an initiator hairpin (H1), the resulting RNA-DNA duplex is thermodynamically more stable than the short native DNA stem—owing to longer complementarity and the intrinsic stability of RNA-DNA helices—thereby driving strand invasion and opening of H1. This conformational change exposes a new initiator sequence that invades an amplifier hairpin (H2), generating a stable H1/H2 duplex while releasing the target RNA to trigger subsequent cycles. Through this turnover, a single RNA molecule catalyzes multiple rounds of strand displacement, leading to exponential signal amplification. In conventional CHA, H1 is labeled with a fluorophore-quencher (F-Q) pair: in the closed state fluorescence is quenched, whereas target-induced opening separates the pair and restores fluorescence, providing a direct readout of the amplification process^22–25^.

By integrating DNA tetrahedra with CHA, we developed a three-dimensional CHA (3D@CHA) nanoplatform that combines structural precision, spatial confinement, and catalytic amplification. In this platform, the tetrahedral scaffold confers high stability, nuclease resistance, efficient cellular uptake, and spatially confined probe positioning, while CHA ensures catalytic turnover and efficient strand displacement. Together, these features overcome the long-standing trade-offs of existing methods by enabling live-cell RNA imaging that is simultaneously stable, noninvasive, highly sensitive, and specific. In addition, spatial confinement of hairpins minimizes background, catalytic cycling generates sustained signal output, and modularity guarantees robustness across diverse cellular contexts, thereby uniting sensitivity, specificity, durability, and biological fidelity within a single platform.

To validate these advantages in practice, we constructed *Ins1*-3D@CHA and *Ins2*-3D@CHA probes. We selected the *Ins1* and *Ins2* mRNAs as ideal model targets because they encode the two insulin isoforms in mice and share a high degree of sequence homology, providing a stringent test for both the sensitivity and discrimination capability of the probes. Using these probes, we precisely distinguished the two highly homologous transcripts and captured mRNA redistribution during metabolic transitions and intercellular transfer, underscoring the platform’s robustness in live-cell applications.

Building on this foundation, we further developed a Förster resonance energy transfer (FRET)-enabled version of the 3D@CHA platform by introducing a single-base recognition site into the H1 hairpin, enabling energy transfer-mediated detection of single-nucleotide variations. A single *KRAS*-3D@CHA probe, derived from this platform, achieved single-base resolution to distinguish wild-type from the clinically relevant *KRAS^G12D^* mutant transcripts, demonstrating its potential for precision molecular diagnostics. The CY3-CY5 readout enabled clear and reliable discrimination both *in vitro* and in living cells.

Together, these advances position 3D@CHA as a versatile and universal platform for RNA biology and precision diagnostics, bridging DNA nanotechnology with catalytic amplification to enable highly sensitive, specific, and noninvasive RNA detection across diverse biological contexts.

## Results

### 1. Design and working mechanism of the 3D@CHA platform

First, based on the sequence characteristics of the target RNA, two distinct and non-overlapping functional regions were selected for probe design (Figure 1A). A longer fragment (∼32 nt, shown in red) was selected as the anchoring domain (AN), which is specifically hybridized and anchored by the probe to ensure stable target capture; a shorter fragment (∼22 nt, shown in green) was defined as the amplification domain (AM), responsible for initiating strand displacement and driving signal amplification. An appropriate spacing was maintained between the two domains to ensure that, once the AN region was anchored to the probe, the AM domain retained sufficient conformational flexibility to freely participate in subsequent amplification reactions.

**Figure 1.**
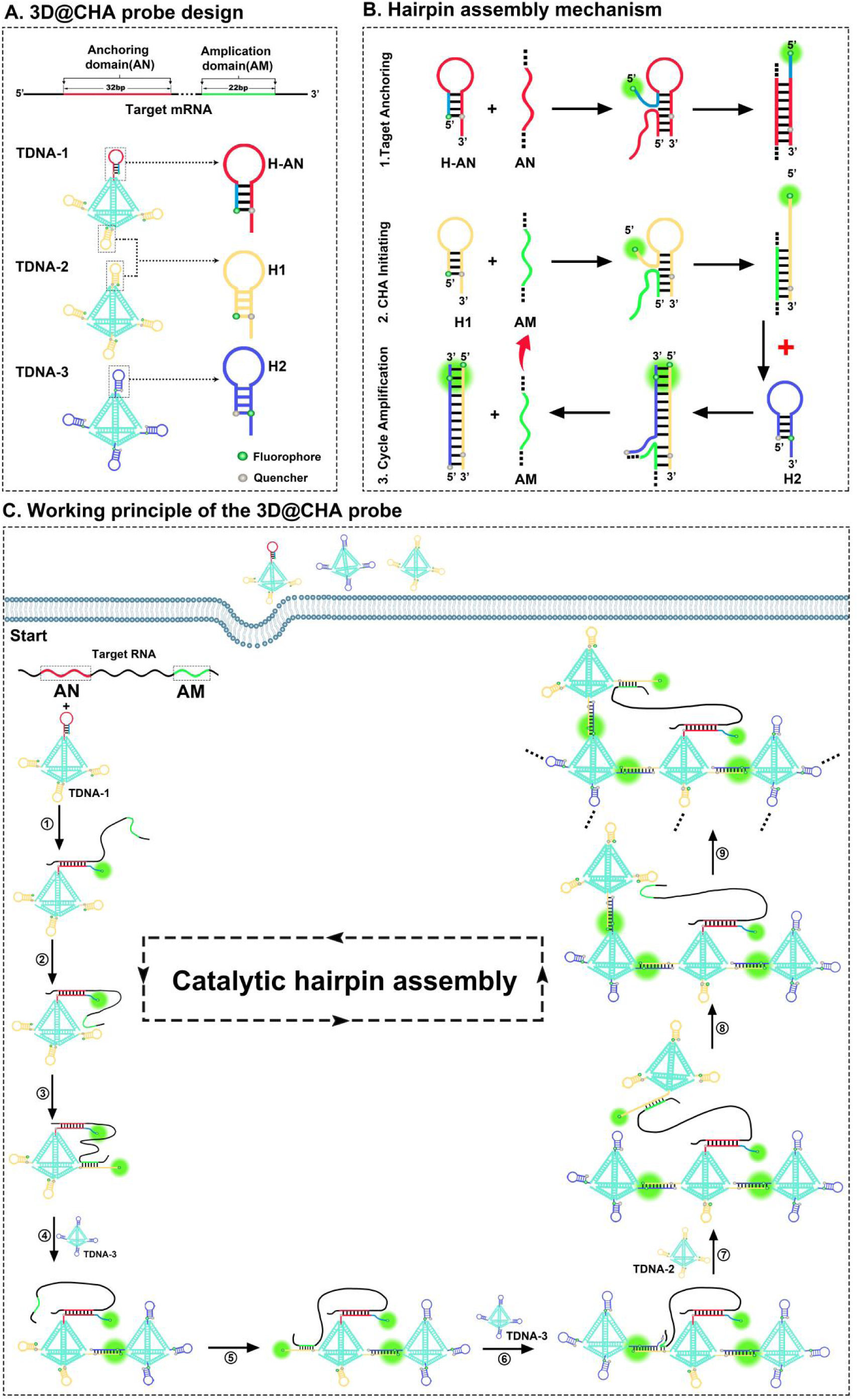
Schematic illustration and working principle of the 3D@CHA probe for RNA imaging. (A) Design and construction of the 3D@CHA probe. The target RNA contains two hybridization regions: an anchoring domain (AN, ∼32 nt, red) and an amplification domain (AM, ∼22 nt, green). The probe comprises three functional tetrahedral DNA modules (TDNA-1, TDNA-2, and TDNA-3). TDNA-1 carries one H-AN (red) and three H1 (yellow) hairpins, TDNA-2 presents four H1 hairpins, and TDNA-3 contains four H2 (blue) hairpins responsible for signal amplification. Fluorophore-quencher (F-Q) pairs are incorporated into H-AN, H1, and H2 to enable fluorescence readout. (B) Reaction mechanism of the hairpin assembly. Step 1. Target anchoring: H-AN (red) hybridizes with the AN (red) region of the target RNA. Step 2. CHA initiation: The AM (green) region of the RNA hybridizes with H1 (yellow), opening the hairpin and triggering CHA. Step 3. Cycle amplification: The released initiator from H1 (yellow) invades H2 (blue), forming H1-H2 (yellow-blue) duplexes that separate the F-Q pairs and restore fluorescence. After each round, the RNA AM strand (green) is displaced and recycled (red arrow), enabling continuous catalytic turnover. (C) Working principle of the 3D@CHA probe. ① TDNA-1 captures the target RNA (red) through hybridization between H-AN and the AN region of the RNA. ② Owing to proximity effects, the amplification domain (AM, green) of the RNA is positioned close to H1 on TDNA-1. ③ The AM hybridizes with H1 (yellow, on TDNA-1), opening its hairpin structure and triggering the CHA reaction. ④ The opened H1 on TDNA-1 hybridizes with H2 on TDNA-3 (blue), separating the fluorophore-quencher (F-Q) pair and releasing the AM strand of the RNA. ⑤ The released AM, guided by spatial proximity, binds to another H1 on TDNA-1, initiating a new round of amplification. ⑥ The newly opened H1 again hybridizes with H2 on TDNA-3, restoring fluorescence and regenerating AM. ⑦ - ⑨ The regenerated AM subsequently hybridizes with H1 on TDNA-2, which carries multiple initiator hairpins, and further propagates across neighboring tetrahedra to drive a cascading amplification reaction.

On this basis, three tetrahedral DNA modules carrying distinct hairpin (stem-loop) structures (TDNA-1, TDNA-2, and TDNA-3) were designed to collectively assemble the nanoscale 3D@CHA probe capable of target recognition and catalytic amplification (Figure 1A). Each TDNA consists of a DNA tetrahedral scaffold and four stem-loop structures positioned at its vertices: TDNA-1 contains one H-AN (red) and three H1 (yellow), enabling specific recognition and anchoring of the target RNA; TDNA-2 carries four H1, serving as a reservoir of initiators to sustain amplification; and TDNA-3 incorporates four H2 (blue) that function as signal amplifiers.

Each hairpin is labeled with a F-Q pair. In the closed state, fluorescence is efficiently quenched, whereas hybridization-induced opening of the stem-loop separates the F-Q pair, leading to fluorescence recovery. Thus, conformational switching of the hairpins is converted into a measurable fluorescence signal, endowing the entire system with self-reporting capability.

As illustrated in Figure 1B and 1C, the overall reaction proceeds through three stages:

Step 1: Target anchoring. The H-AN on TDNA-1 hybridizes specifically with the AN region of the RNA, anchoring the target molecule onto the nanoscale scaffold (Figure 1B, Step 1; Figure 1C, Step ①).

Step 2: CHA initiation. Due to spatial proximity, the AM region of the immobilized RNA more readily hybridizes with H1 on TDNA-1 (Figure 1C, Step ②), thereby opening the H1 and initiating CHA reaction (Figure 1B, Step 2; Figure 1C, Step ③). At this stage, the conformational transition of H1 separates the F-Q pair and restores fluorescence.

Step 3: Cyclic amplification. The opened H1 further hybridizes with H2 on TDNA-3 (Figure 1B, Step 3; Figure 1C, Step ④), separating another F-Q pair and producing an additional fluorescence signal while releasing the AM strand (Figure 1B, red arrow). The released AM can rehybridize with another H1 on TDNA-1 (Figure 1C, Step ⑤) or randomly collide with H1 on TDNA-2 (Figure 1C, Step ⑦), reopening hairpins and driving new rounds of H1-H2 hybridization (Figure 1C, Steps ⑥ and ⑧). This turnover continuously regenerates AM and results in iterative fluorescence amplification (Figure 1C, Step ⑨).

Through this cascade reaction—governed by hairpin dynamics and spatially confined within the tetrahedral architecture—the 3D@CHA probe achieves autonomous cyclic CHA amplification and pronounced signal enhancement (Figure 1C). In other words, the tetrahedron serves as the spatial carrier, while the hairpins act as the reactive units; together, they constitute a nanoscale, self-driven three-dimensional amplification network. A single RNA molecule can thereby trigger multiple amplification cycles, generating an intense and spatially restricted fluorescence signal suitable for *in situ* RNA imaging.

The complete working process of the 3D@CHA probe— including target anchoring, hairpin opening, cyclic amplification, and fluorescence generation—was dynamically visualized, as shown in Video 1.

### 2. Construction and validation of the 3D@CHA probe

To validate the feasibility of the 3D@CHA probe, we first examined its sequence design and hierarchical assembly. Figure S1 illustrates the stepwise construction process, in which single-stranded DNAs (S1-S9, Figure S1A) fold into defined stem-loop structures and subsequently assemble into triangular faces (Figure S1B) and tetrahedral units(Figure S1C), ultimately forming three functional TDNA modules (TDNA-1, TDNA-2, and TDNA-3) that together constitute the complete probe (Figure S1D).

After sequence design, the structural assembly of the probe was experimentally confirmed. As shown in Figure 2A, the electrophoretic bands gradually shifted to lower mobility with the sequential addition of S1, S3, S4, and S5, revealing a progressive increase in molecular size and an orderly formation of the tetrahedral structure. This result specifically demonstrated the successful assembly of TDNA-1, which integrates one H-AN and three H1 to form a well-defined tetrahedral module.

**Figure 2.**
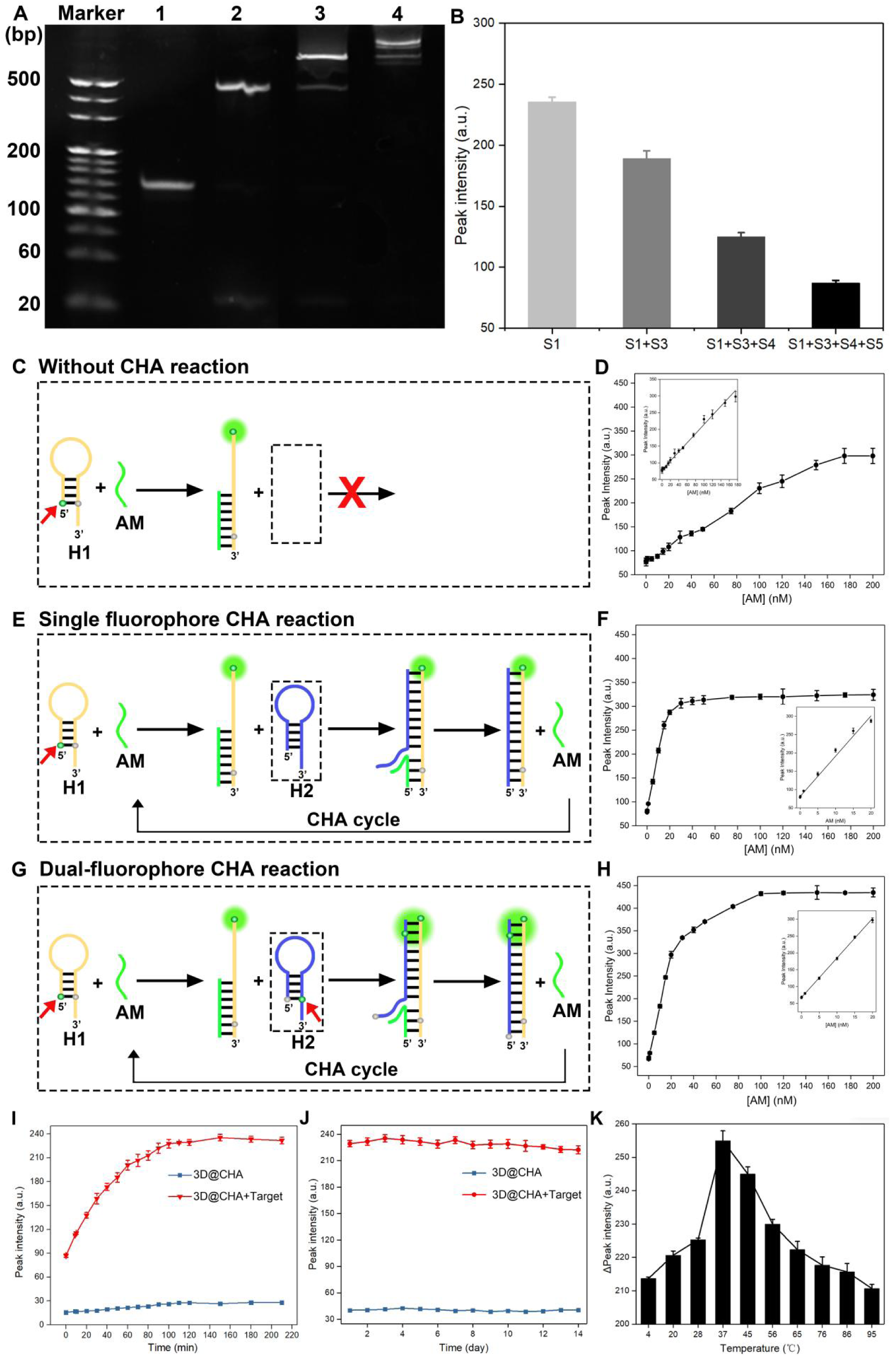
Construction and evaluation of the 3D@CHA signal amplification strategy in solution. (A) Polyacrylamide gel electrophoresis (PAGE) analysis confirming the stepwise formation of the TDAN1. Lane 1: S1; Lane 2: S1+S3; Lane 3: S1+S3+S4; Lane 4: S1+S3+S4+S5. (B) Fluorescence responses corresponding to each assembly stage in (A), showing progressive fluorescence quenching upon tetrahedron formation. (C) Schematic illustration of the detection system without CHA amplification. The red cross indicates that no CHA reaction occurs, and the red arrows mark the positions of the F-Q pairs on the stem-loop structures. (D) Fluorescence response curves of the non-amplified system containing AM and F-Q-labeled H1 at different AM concentrations (0-200 nM). (E) Schematic illustration of the single-fluorophore CHA reaction system. (F) Fluorescence response of the single-fluorophore CHA system at different AM concentrations (0-200 nM). (G) Schematic illustration of the dual-fluorophore CHA reaction system. (H) Fluorescence response of the dual-fluorophore CHA system at different AM concentrations (0-200 nM). (I) Real-time monitoring of fluorescence intensity of the 3D@CHA probe in the presence or absence of target RNA (AN+AM), showing fluorescence signal changes over 220 min and reaching saturation at 120 min. (J) Fluorescence stability of the 3D@CHA probe after 14 days of storage, with or without target RNA (AN+AM). (K) Temperature-dependent fluorescence intensity difference between target and non-target systems, showing that the fluorescence signal was strongest at 37 °C.

Fluorescence analysis further supported these results (Figure 2B). In the single-stranded state, the F-Q pairs were spatially separated, producing strong fluorescence, whereas tetrahedral assembly brought them into close proximity, resulting in effective fluorescence quenching. Collectively, these results verify the successful and ordered construction of the designed DNA tetrahedral probe and establish a structural foundation for subsequent amplification assays.

We next evaluated the amplification efficiency of the 3D@CHA probe under different configurations to identify the optimal amplification strategy (Figure 2C-H). In the absence of H2 (Figure 2C), no CHA reaction occurred; the system containing only AM and H1 (F-Q labeled) showed a gradual fluorescence increase with target concentration but remained unsaturated within 0-200 nM (Figure 2D), corresponding to a relatively high limit of detection (LOD). Introducing unlabeled H2 initiated the CHA cycle (Figure 2E-F), enhancing fluorescence intensity by approximately 5-fold and reducing the LOD from 1.0 nM to 0.209 nM (Table 1), indicating improved amplification efficiency.

**Table. 1.** Detection performance of amplification strategy.

| Sample | linear relationship<br>(nM) | Linear equation | LOD (nM) |
| --- | --- | --- | --- |
| AM | 0-175 | $y = 1.360x + 78.515$<br>($n = 3, R^2 = 0.995$) | 1.0 |
| AM | 0-20 | $y = 11.005x + 80.965$<br>($n = 3, R^2 = 0.987$) | 0.209 |
| AM | 0-20 | $y = 11.763x + 67.436$<br>($n = 3, R^2 = 0.999$) | 0.01 |

Encouraged by this improvement, we further sought to enhance signal output by introducing an additional F-Q pair on H2, constructing a dual-fluorophore configuration (Figure 2G-H). This dual-labeled CHA system produced the strongest fluorescence response and achieved an LOD of 0.01 nM — approximately 20-fold higher sensitivity than the single-fluorophore CHA and two orders of magnitude greater than the non-amplified system. These results demonstrate that incorporating two F-Q pairs markedly improves both signal gain and detection sensitivity, making the dual-fluorophore 3D@CHA design the most efficient amplification strategy.

We further characterized the reaction kinetics and stability of the optimized 3D@CHA probe. As shown in Figure 2I, fluorescence intensity gradually increased in the presence of target AM and AN sequences and reached saturation at approximately 120 min, indicating completion of the strand-displacement reaction within 2 h; thus, 2 h was defined as the optimal reaction time. Under these conditions, fluorescence signals remained stable for 14 days in both target and non-target systems, exhibiting minimal fluctuation and no observable decay (Figure 2J), confirming the excellent photochemical and structural stability of the probe. Temperature-dependent analysis (Figure 2K) showed that fluorescence intensity peaked at 37 ℃ and declined at higher temperatures, demonstrating that the probe performs optimally under physiological conditions and is well suited for cellular and *in vivo* RNA detection.

Collectively, these results confirm the precise structural formation, robust amplification capability, and exceptional stability of the 3D@CHA probe, providing a solid foundation for its application in *in situ* RNA imaging.

### 3. Sensitive and specific detection of *Ins1* and *Ins2* mRNA *in vitro*

To evaluate the sensitivity and specificity of the 3D@CHA probe for RNA detection, two highly homologous insulin transcripts— *Ins1* and *Ins2* —were selected as model targets. Corresponding FAM-labeled *Ins1*-3D@CHA and CY5-labeled *Ins2*-3D@CHA probes were constructed for dual-fluorescence analysis.

As shown in Figure 3A (*Ins1*-3D@CHA) and Figure 3G (*Ins2*-3D@CHA), in the absence of target RNA, the stem-loop structures remained intact and fluorescence was efficiently quenched. Addition of the AN fragment partially restored fluorescence, while subsequent introduction of the AM fragment triggered strand-displacement amplification and markedly enhanced the signal. When both AN and AM were present, fluorescence intensity reached its maximum, confirming that AM acts as the primary initiator and that AN-AM cooperation synergistically amplifies the response.

**Figure 3.**
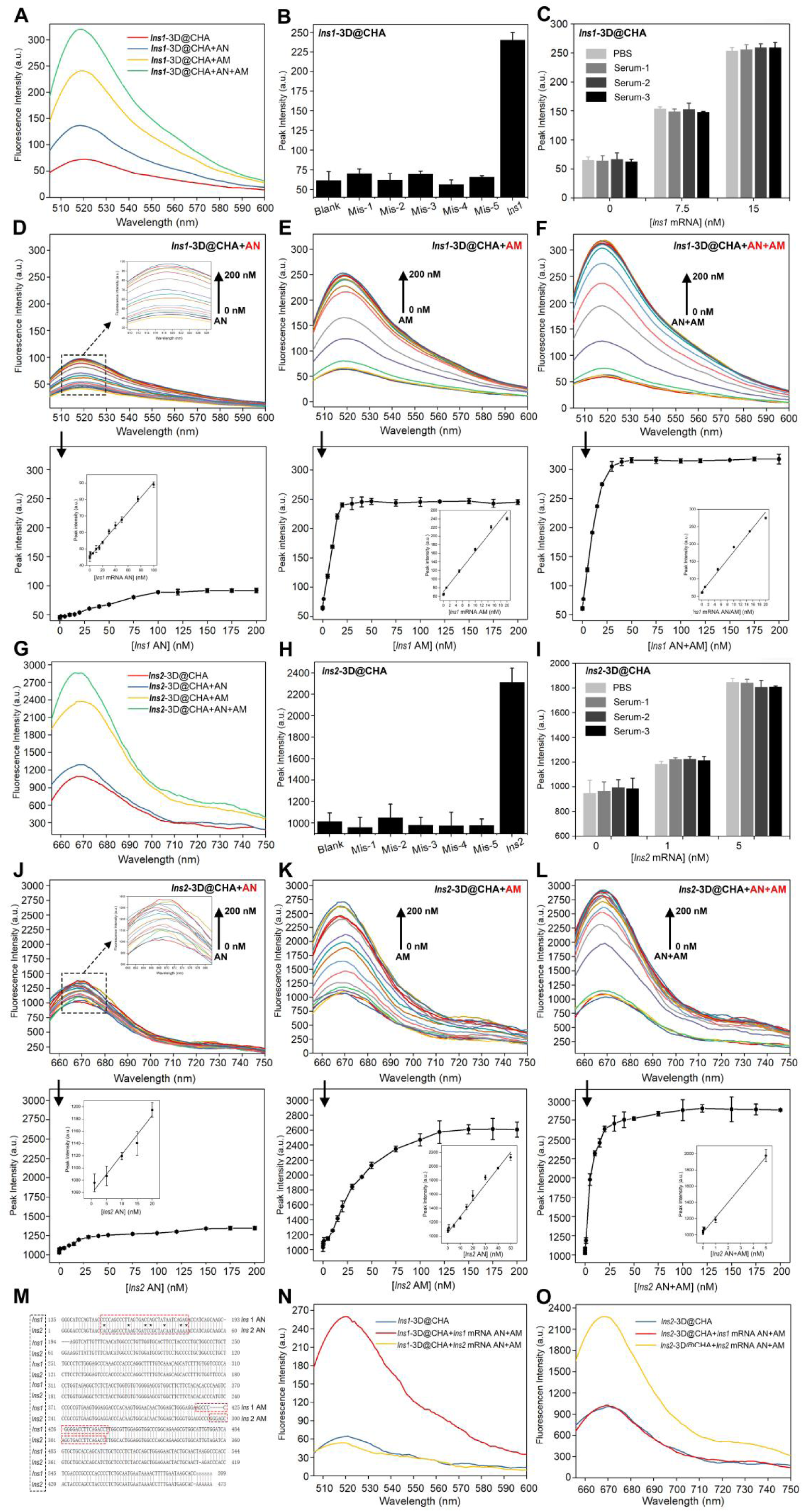
Evaluation of the *Ins1*- and *Ins2*-3D@CHA probes in solution. (A) Fluorescence spectra of the *Ins1*-3D@CHA probe responding to the *Ins1* mRNA-derived AN, AM, and AN+AM fragments. (B) Fluorescence intensities of the *Ins1*-3D@CHA probe toward the fully matched *Ins1* target (AN+AM) and five mismatched sequences, each containing three randomly distributed base substitutions. (C) Detection of *Ins1* mRNA (AN+AM) in three independently collected serum samples using the *Ins1*-3D@CHA probe, performed with three different target concentrations. (D-F) Fluorescence spectra of the *Ins1*-3D@CHA probe with increasing concentrations (0-200 nM) of (D) *Ins1* AN, (E) *Ins1* AM, and (F) *Ins1* AN+AM fragments (upper). Lower panels: calibration curves correlating fluorescence peak intensity with target concentration; insets show the linear fitting ranges. (G) Fluorescence spectra of the *Ins2*-3D@CHA probe responding to the *Ins2* mRNA-derived AN, AM, and AN+AM fragments. (H) Fluorescence intensities of the *Ins2*-3D@CHA probe toward the fully matched *Ins2* target (AN+AM) and five mismatched sequences (each containing three base mismatches). (I) Detection of *Ins2* mRNA in three independent serum samples using the *Ins2*-3D@CHA probe at three different target concentrations. (J-L) Fluorescence spectra of the *Ins2*-3D@CHA probe with increasing concentrations (0-200 nM) of (J) *Ins2* AN, (K) *Ins2* AM, and (L) *Ins2* AN+AM fragments (upper). Lower panels: calibration curves between fluorescence peak intensity and target concentration; insets show the corresponding linear fitting ranges. (M) Sequence alignment of *Ins1* and *Ins2* mRNAs, highlighting regions with the largest nucleotide differences (boxed in red) selected for probe design. (N-O) Fluorescence responses of the *Ins1*-3D@CHA (N) and *Ins2*-3D@CHA (O) probes toward *Ins1* mRNA (AN+AM), *Ins2* mRNA (AN+AM), and blank controls, demonstrating high target selectivity and absence of cross-interference.

Fluorescence spectra were recorded across gradient concentrations of target fragments to generate calibration curves and evaluate quantitative performance (upper panels of Figure 3D-3F, 3J-3L; Table 2 and 3). LOD was determined using the 3σ method (lower panels of Figure 3D-3F, 3J-3L).

**Table. 2.** Detection performance of *Ins1*-3D@CHA.

| Sample | linear relationship<br>(nM) | Linear equation | LOD (nM) |
| --- | --- | --- | --- |
| *Ins1* mRNA AN | 0-100 | $y = 0.44954x + 45.342$<br>($n = 3, R^2 = 0.993$) | 1.0 |
| *Ins1* mRNA AM | 0-20 | $y = 9.3278x + 70.672$<br>($n = 3, R^2 = 0.972$) | 0.117 |
| *Ins1* mRNA AN+AM | 0-20 | $y = 11.390x + 64.336$<br>($n = 3, R^2 = 0.989$) | 0.01 |

**Table. 3.** Detection performance of *Ins2*-3D@CHA.

| Sample | linear relationship<br>(nM) | Linear equation | LOD (nM) |
| --- | --- | --- | --- |
| *Ins2* mRNA AN | 0-20 | $y = 6.404x + 1057.738$<br>($n = 3, R^2 = 0.965$) | 1.0 |
| *Ins2* mRNA AM | 0-50 | $y = 6.404x + 1057.738$<br>($n = 3, R^2 = 0.965$) | 0.106 |
| *Ins2* mRNA AN+AM | 0-5 | $y = 186.319x + 1028.819$<br>($n = 3, R^2 = 0.986$) | 0.0085 |

\*in which $y$ stands for the peak intensity and $x$ is sample concentration.

For *Ins1*-3D@CHA (Figure 3D-3F), fluorescence spectra exhibited gradual signal enhancement with increasing concentrations of AN (Figure 3D), AM (Figure 3E), and AN+AM (Figure 3F) fragments. In our probe design, the H-AN strand was also modified with a F-Q pair; therefore, hybridization with the target AN region produced a measurable fluorescence signal, indicating partial strand opening. Addition of the AM fragment initiated catalytic hairpin assembly and further enhanced fluorescence, while the co-presence of AN and AM yielded the strongest signal, demonstrating that cooperative activation between the two domains effectively sustains the CHA reaction. The calculated LOD values (1.0 nM for AN, 0.117 nM for AM, and 0.01 nM for AN+AM) confirmed the robustness and efficiency of this amplification process.

Similarly, for *Ins2*-3D@CHA (Figure 3J-3L), fluorescence intensity progressively increased under the AN, AM, and AN+AM conditions. Consistent with the *Ins1*-3D@CHA results, the AM fragment significantly enhanced fluorescence, and the combination of AN and AM generated the highest signal, confirming that the cooperative amplification mechanism operates reproducibly in both probe systems. The corresponding LOD values (1.0 nM for AN, 0.106 nM for AM, and 0.0085 nM for AN+AM) further verified the reliability and amplification efficiency of the 3D@CHA-based detection strategy.

To further assess sequence specificity, five RNA fragments containing three mismatched bases at random positions were designed as negative controls. For both *Ins1*-3D@CHA (Figure 3B) and *Ins2*-3D@CHA (Figure 3H), all mismatched sequences failed to trigger hairpin opening and produced negligible fluorescence, whereas fully complementary targets generated strong signals, indicating high sequence discrimination and excellent specificity.

In addition, to evaluate probe stability and reliability in complex biological matrices, target RNAs were spiked at varying concentrations into three independent serum samples. As shown in Figure 3C and 3I, fluorescence intensity increased proportionally with target concentration, and no significant variations were observed among different serum samples. These results confirm that the 3D@CHA probe maintains consistent amplification performance and stability even under complex physiological conditions.

Following systematic validation of the *Ins1*-3D@CHA and *Ins2*-3D@CHA probes (Figure 3A-3L), which confirmed their excellent amplification efficiency and reliable detection performance, the deeper purpose of selecting *Ins1* and *Ins2* as targets was to further explore whether the 3D@CHA system could precisely discriminate RNA sequences with extremely high similarity. As shown in Figure 3M, *Ins1* and *Ins2* mRNAs share approximately 90% sequence identity. The regions with the most distinct nucleotide differences (highlighted in red) were selected for probe design to maximize sequence discrimination.

To verify this, the *Ins1*-3D@CHA and *Ins2*-3D@CHA probes were individually tested for cross-reactivity. As illustrated in Figure 3N, the *Ins1*-3D@CHA probe produced a strong fluorescence signal only in the presence of *Ins1* mRNA, whereas negligible fluorescence was observed for *Ins2* mRNA or the blank control. Conversely, the *Ins2*-3D@CHA probe (Figure 3O) yielded a distinct CY5 fluorescence signal exclusively upon addition of *Ins2* mRNA, with no detectable response to *Ins1* mRNA or in the absence of target. These findings indicate that both probes specifically recognize their corresponding mRNA targets, demonstrating that the 3D@CHA system can accurately distinguish highly homologous RNA sequences with excellent specificity and without cross-interference.

Such a high degree of sequence selectivity, coupled with the intrinsic stability of the tetrahedral framework, strongly underscores the system’s potential for *in situ* and *in vivo* RNA detection applications.

### 4. Discrimination and dynamic imaging of insulin mRNAs in living cells

Following *in vitro* validation of the 3D@CHA probe’s high sensitivity and specificity for highly homologous *Ins1* and *Ins2* mRNAs, we further evaluated its accuracy in target recognition and imaging performance in living cells.

Human embryonic kidney cells (293T), 293T cells overexpressing *Ins1* (293T*^Ins1+^*) and *Ins2* (293T*_Ins2_*_+_) (overexpression verified in Figure S2), and the mouse pancreatic β-cell line MIN6 (mouse insulinoma MIN6 cells) were used as model systems.

Before live-cell experiments, the biocompatibility and cytotoxicity of the DNA tetrahedral probes were examined. CCK-8 assays showed that probe treatment did not affect 293T cell viability after incubation for 0-9 h at 100 nM (Figure S3A) or across 0-100 nM concentrations for 12 h (Figure S3B), confirming good cellular safety and compatibility.

We next investigated the probes’ target recognition and intracellular detection performance across different cell models. In 293T*^Ins2+^* cells, incubation with the *Ins2*-3D@CHA probe resulted in a time-dependent fluorescence increase over 12 h (Figure 4A), demonstrating efficient cellular uptake, high intracellular stability, and sustained detection capability.

**Figure 4.**
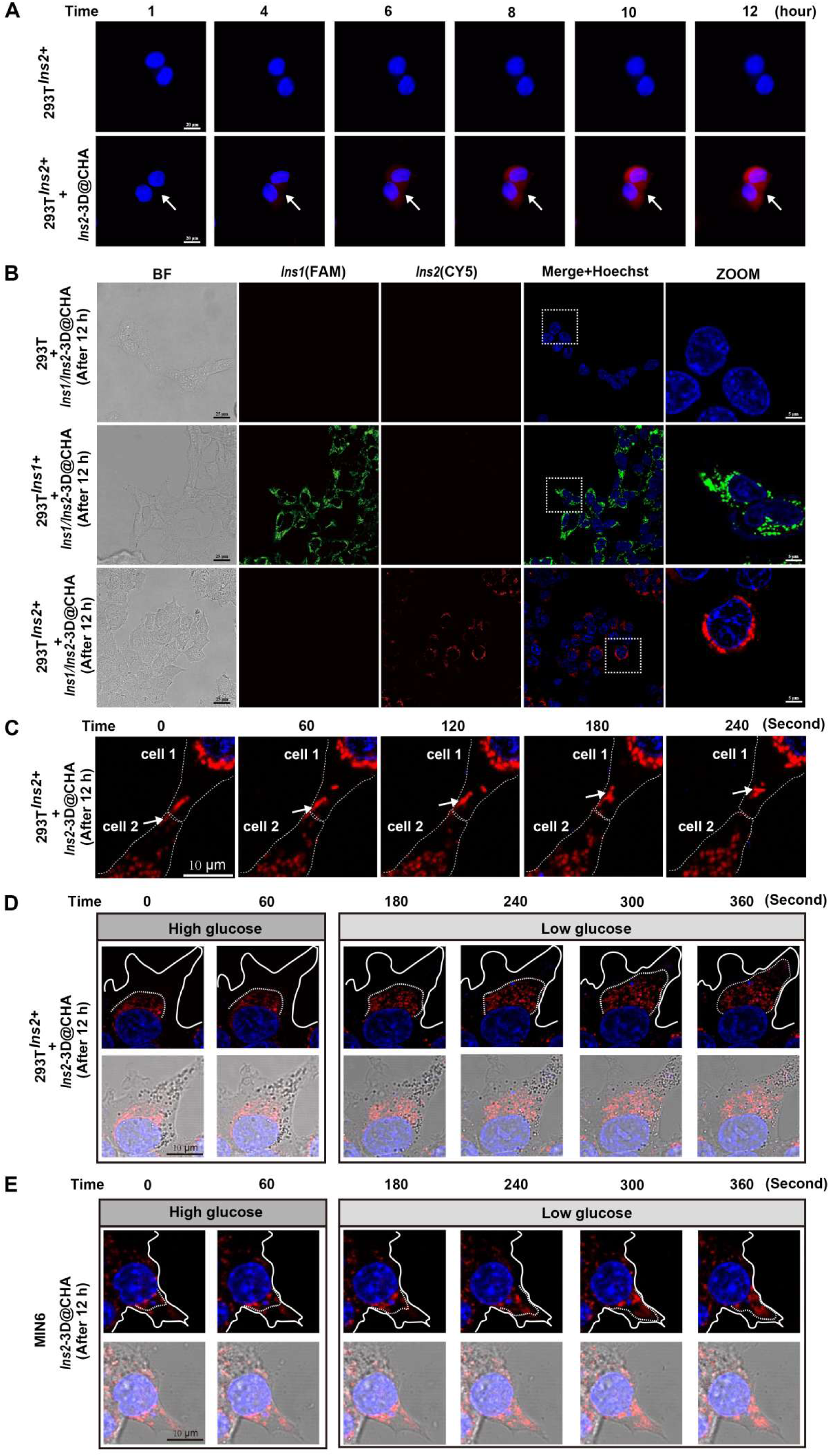
3D@CHA platform-mediated imaging of mRNA in living cells. (A) Time-lapse confocal fluorescence images of 293T*^Ins2+^* cells incubated with or without the Ins2-3D@CHA probe for different time periods (1, 4, 6, 8, 10, and 12 h). Red fluorescence represents CY5 signals from hybridized probes, and blue fluorescence represents Hoechst-stained nuclei. Scale bar, 20 μm. (B) Confocal fluorescence imaging of 293T, 293T*^Ins1+^*, and 293T*^Ins2+^* cells co-incubated with *Ins1*-3D@CHA (FAM channel, green) and *Ins2*-3D@CHA (CY5 channel, red) probes for 12 h. Merged images include Hoechst-stained nuclei (blue), and ZOOM panels display magnified regions of interest. BF, bright-field images. Scale bars, 25 μm. (C) Real-time confocal imaging of *Ins2* mRNA trafficking between adjacent 293T*^Ins2+^* cells after 12 h incubation with the *Ins2*-3D@CHA probe. Dashed white lines outline cell boundaries. Sequential frames (0-240 s) show the gradual transfer of fluorescent mRNA signal from cell 2 to cell 1. Scale bar, 10 μm. (D) Dynamic redistribution of *Ins2* mRNA in 293T*^Ins2+^* cells during a glucose-switch experiment. Cells were pre-incubated with the *Ins2*-3D@CHA probe for 12 h under high-glucose (4.5 g/L) conditions and then switched to low-glucose (1.0 g/L) medium. Sequential images (0-360 s) reveal that the *Ins2* mRNA signal (red) initially concentrated around the nucleus gradually diffused into the cytoplasm. Upper panels show fluorescence channels; lower panels display merged fluorescence with bright-field images. Scale bar, 10 μm. (E) Time-lapse imaging of glucose-responsive mRNA dynamics in MIN6 β-cells. After 12 h incubation with the *Ins2*-3D@CHA probe, a medium change from high-glucose (4.5 g/L) to low-glucose (1.0 g/L) induced redistribution of *Ins2* mRNA (0-360 s). Upper panels show fluorescence channels; lower panels display merged fluorescence and bright-field images. Scale bar, 10 μm.

Furthermore, both *Ins1*-3D@CHA and *Ins2*-3D@CHA probes were simultaneously introduced into the same cell system to evaluate their orthogonal detection capability. After 12 h of incubation, no fluorescence was observed in wild-type 293T cells, whereas distinct FAM and CY5 signals appeared exclusively in 293T*^Ins1+^* and 293T*^Ins2+^* cells, respectively (Figure 4B). These findings confirm that the 3D@CHA probes can accurately discriminate highly homologous transcripts at the cellular level. Similarly, in MIN6 β-cells, co-incubation with both probes (100 nM each, 12 h) led to time-dependent increase in fluorescence (Figure S4), indicating high cell permeability and structural stability for synchronous dual-target imaging.

Accordingly, we confirmed that under live-cell detection conditions, the probe fully hybridized with its target within 12 hours, reaching a stable fluorescence signal. Based on this, we further monitored the cells and found that the probe could sensitively track dynamic mRNA behavior in living systems.

In 293T*^Ins2+^* cells, the *Ins2*-3D@CHA probe clearly revealed the transfer of *Ins2* mRNA between adjacent cells (Figure 4C, Video 2), indicating its capability for real-time monitoring of intercellular mRNA trafficking. Moreover, in both 293T*^Ins2+^* (Figure 4D) and MIN6 (Figure 4E) cells, when the culture conditions were switched from high to low glucose, the *Ins2* mRNA signal redistributed from perinuclear enrichment to cytoplasmic diffusion (Videos 3-4), further demonstrating that the probe enables stable imaging and dynamic tracking of mRNA spatial redistribution under different cell types and environmental contexts.

### 5. Improving single-base mutation discrimination through H1 structural recognition and FRET-based readout enhancement

The *KRAS^G12D^* mutation—a single-nucleotide substitution at codon 12 (GGT→GAT) that replaces glycine with aspartic acid—is one of the most prevalent oncogenic drivers in pancreatic, colorectal, and lung cancers. Sensitive and accurate detection of this mutation is therefore essential for early diagnosis, therapeutic monitoring, and the development of precision-targeted therapies. To establish a proof-of-concept system capable of discriminating single-base mutations at the RNA level, we selected *KRAS^G12D^* as a representative model.

We first selected the H1 hairpin as the mutation recognition unit within the CHA circuit, given its dual role as both the amplification core and the rate-limiting element in strand displacement. The mutation site was embedded within the stem region of H1, allowing single-base mismatches to directly interfere with hairpin opening and thereby modulate the overall amplification efficiency.

To maximize mismatch sensitivity, we systematically evaluated the impact of recognition-site positioning within the H1 stem. Three potential sites were designed: the hairpin initiation region (site I), middle region (site II), and terminal region (site III) (Figure 5A). Because the energy barrier for strand displacement is mainly determined by the local base-pair stability near the hairpin opening site, a mismatch closer to the initiation region is theoretically expected to cause greater structural destabilization and stronger inhibition of hairpin opening. Consistent with this prediction, experimental results and thermodynamic simulations showed a high degree of agreement: Site I (initiation region) produced the most pronounced fluorescence difference between wild-type and mutant *KRAS*, whereas sites II and III exhibited markedly reduced discrimination efficiency (Figure S5-S7). Complementary NUPACK analyses further confirmed that site I possessed the lowest free energy (ΔG) and smallest energy barrier, resulting in the highest mismatch sensitivity (Figure S8-S10). Accordingly, site I was identified as the optimal recognition position and adopted for all subsequent experiments.

**Figure 5.**
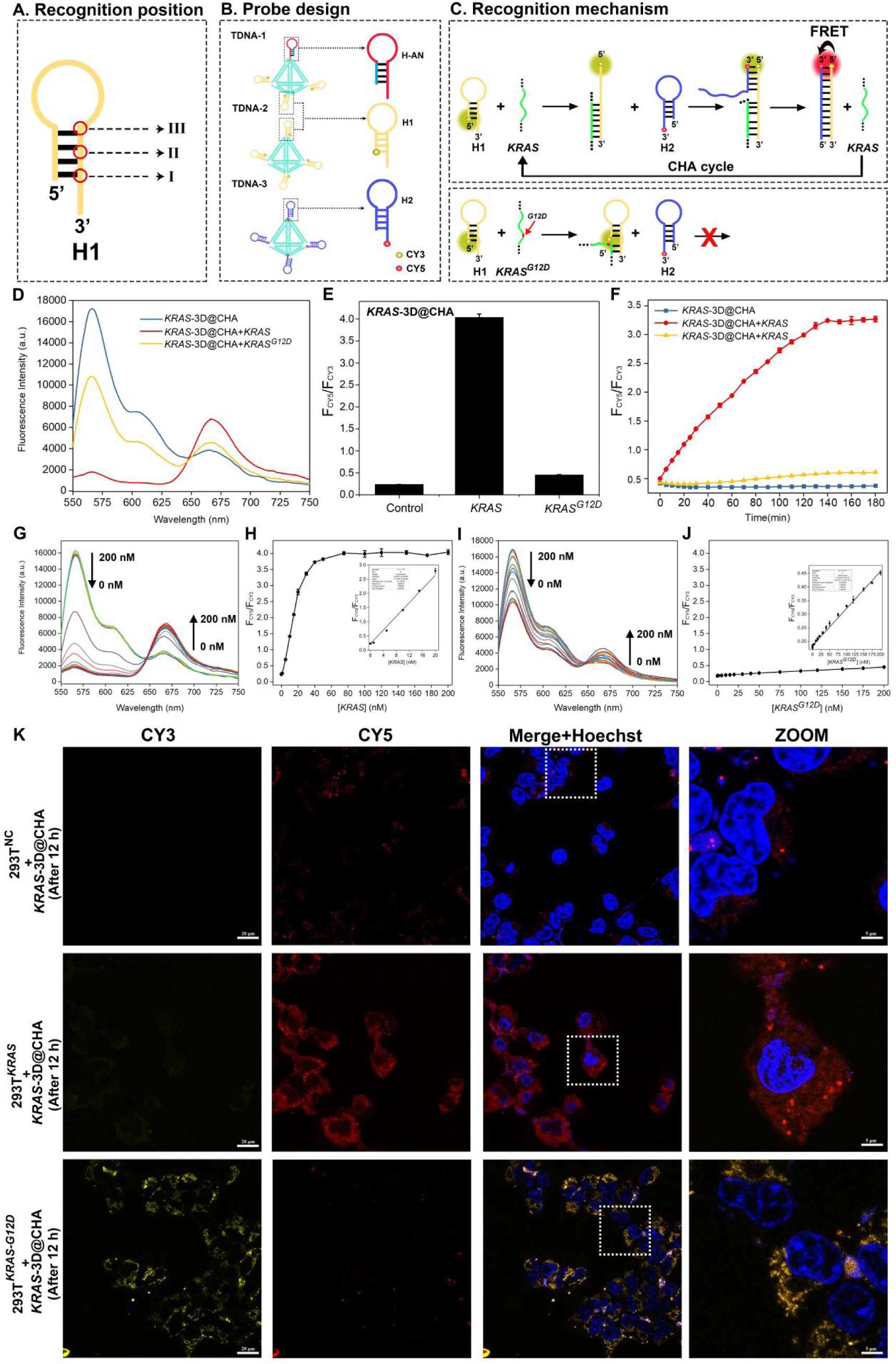
Imaging and discrimination of single-base mutation mRNA via the FRET-enhanced 3D@CHA amplification system. (A) Optimization of the mutation recognition site within the H1 hairpin. Three potential recognition positions were tested—site I (initiation region), site II (middle), and site III (terminal region). (B) Schematic illustration of the construction of functional probes by integrating three hairpins (H-AN, H1, and H2) within the 3D@CHA framework. The donor fluorophore CY3 was conjugated to the 5’ end of H1, and the acceptor fluorophore CY5 was attached to the 3’ end of H2 to form a FRET pair. (C) Working mechanism of the 3D@CHA platform for distinguishing *KRAS* and *KRAS^G12D^* mRNAs. Upper panel: Perfectly matched *KRAS* triggers strand displacement and the catalytic hairpin assembly (CHA) reaction between H1 and H2, bringing CY3 (donor) and CY5 (acceptor) into close proximity and generating strong FRET signals. The target *KRAS* is released after each cycle, enabling continuous amplification and cyclic detection. Lower panel: In contrast, the single-base mismatch in *KRAS^G12D^* hinders strand displacement and prevents effective H1-H2 hybridization, resulting in a blocked CHA cycle (red cross) and a markedly reduced FRET response. (D) Fluorescence spectra of the 3D@CHA system after addition of *KRAS* and *KRAS^G12D^* mRNAs. (E) Histogram of the fluorescence intensity ratio (F_CY5_/F_CY3_) of the 3D@CHA system in the presence of *KRAS* and *KRAS^G12D^* mRNAs. (F) Real-time monitoring of the fluorescence response of the *KRAS*-3D@CHA system toward *KRAS*, *KRAS^G12D^*, and blank (no-target control) samples. (G) Fluorescence spectra of the 3D@CHA system upon addition of *KRAS* mRNA at varying concentrations (0-200 nM). (H) Calibration curve of F_CY5_/F_CY3_ versus *KRAS* mRNA concentration; the inset displays the linear correlation between F_CY5_/F_CY3_ and concentration in the low-nanomolar range. (I) Fluorescence spectra of the system in response to increasing concentrations (0-200 nM) of mutant *KRAS^G12D^* mRNA. (J) Calibration curve of F_CY5_/F_CY3_ versus *KRAS^G12D^* mRNA concentration; the inset shows the linear fit of F_CY5_/F_CY3_ values with consistently low ratios across all samples. (K) Confocal laser scanning microscopy images of the 3D@CHA imaging probes co-incubated with 293T^NC^, 293T*^KRAS^*, and 293T*^KRAS-G12D^* cells for 12 h. CY3 (yellow) represents the donor fluorophore on H1, CY5 (red) represents the acceptor fluorophore on H2, and Hoechst (blue) stains cell nuclei. Scale bars, 20 μm.

Based on our previously developed 3D@CHA platform, we next designed a *KRAS*-3D@CHA probe (Figure S11) incorporating the optimized H1 recognition site. In this configuration, the probe consists of two hairpin strands—H1 (yellow) and H2 (blue)—embedded within a three-dimensional tetrahedral DNA framework. Upon hybridization with the target *KRAS* mRNA, the 5’ region of H1 binds to the complementary sequence, inducing hairpin opening and exposing a toehold for strand displacement with H2. The reaction generates an H1-H2 duplex and releases the *KRAS* trigger to initiate additional CHA cycles, thereby producing a strong fluorescence “turn-on” signal (Figure S11A). Although the fluorescence intensity differed between wild-type and mutant *KRAS* samples, the difference in signal strength within a single detection channel was insufficient for clear visual discrimination, limiting the ability to reliably distinguish between the two forms (Figure S11B-11D).

To overcome this limitation and enhance signal discernibility, we introduced a FRET-based ratiometric readout to provide a spectrally distinct and internally calibrated signal output (Figure 5B). In this configuration, CY3 and CY5 were employed as the donor-acceptor fluorophore pair, labeled respectively at the 5’ end of H1 and the 3’ end of H2, while H-AN served as an anchoring domain without fluorescence contribution. For wild-type *KRAS* mRNA (Figure 5C, upper), efficient strand displacement brought CY3 and CY5 into close proximity, resulting in strong CY5 fluorescence enhancement and CY3 quenching through FRET. In contrast, for the *KRAS^G12D^* mutant mRNA (Figure 5C, lower), the single-base mismatch disrupted H1-H2 hybridization, blocking the amplification reaction (indicated by the red cross) and thereby preventing energy transfer. This produced distinct color-dependent emission differences, with the mutant sample exhibiting markedly reduced FRET intensity.

By integrating H1 structural recognition with FRET-based ratiometric detection, this optimized *KRAS*-3D@CHA system successfully converted subtle hybridization differences into visually and spectrally distinguishable outputs, enabling precise and reliable discrimination of single-base mutations.

*In vitro* experiments demonstrated that wild-type *KRAS* generated strong CY5 fluorescence accompanied by significant CY3 quenching, whereas *KRAS^G12D^* produced only weak CY5 emission and residual CY3 fluorescence (Figure 5D). The FRET efficiency, calculated as the fluorescence intensity ratio (F_CY5_/F_CY3_), was significantly higher for wild-type samples than for mutant samples (Figure 5E). Kinetic analysis revealed that the F_CY5_/F_CY3_ ratio of wild-type *KRAS* rapidly increased within approximately 140 minutes and reached a plateau, while *KRAS^G12D^*maintained low reaction efficiency throughout (Figure 5F), confirming that the platform effectively differentiates reaction kinetics between wild-type and mutant transcripts.

Concentration-gradient experiments further validated the system’s sensitivity and robustness: as the concentration of wild-type *KRAS* transcripts increased, CY5 fluorescence intensified and CY3 emission decreased, with FRET efficiency steadily rising and saturating at approximately 50 nM (Figure 5G-H). In contrast, *KRAS^G12D^* samples exhibited a negligible response across the 0-200 nM range, with weak FRET response (Figure 5I-J), reaffirming the high selectivity for mutation detection.

To verify the probe’s mutation discrimination capability in cellular systems, 293T overexpression models were generated in which cells expressed either the wild-type or mutant form of *KRAS*. Three experimental groups were established: vector control (293T^NC^), *KRAS*-WT (293T*^KRAS^*), and *KRAS-G12D* (293T*^KRAS-G12D^*). Successful overexpression was confirmed by Western blot analysis prior to fluorescence imaging. (Figure S12). Strong CY5 fluorescence with nearly complete CY3 quenching was observed in 293T*^KRAS^* cells, whereas 293T*^KRAS-G12D^* cells displayed weak CY5 and residual CY3 signals. In the negative control 293T^NC^ cells, only weak CY5 fluorescence was detected (Figure 5K). These cellular observations were consistent with the *in vitro* data, confirming that the FRET-enhanced 3D@CHA system enables accurate identification and discrimination of single-base mutant transcripts within living cells.

In summary, by introducing a strategically positioned mutation recognition site within the initiation region of the H1 hairpin and integrating a FRET-based dual-fluorescence readout, the optimized 3D@CHA platform achieves simultaneous enhancement of structural recognition and signal resolution. This system demonstrates outstanding single-base precision both *in vitro* and in living cells, overcoming the limitations of conventional fluorescence probes and providing a powerful and versatile tool for RNA-level mutation analysis and precision medicine research.

## Discussion

Visualizing RNA dynamics in living systems remains highly challenging because of the intrinsic complexity of RNA behavior. Existing imaging methods suffer from low sensitivity, high background, and poor biocompatibility, hindering faithful resolution of endogenous RNA spatiotemporal patterns. Conventional strategies, such as MS2-GFP tagging or molecular beacons, have enabled visualization of specific transcripts but often require genetic modification or exhibit high background fluorescence, limiting their applicability for endogenous RNA imaging ^26–28^. Achieving single-base precision in living cells further demands exceptional probe stability and discrimination accuracy and remains a long-standing challenge in molecular detection.

To address these challenges, we developed an integrated 3D@CHA nanoplatform that enables real-time imaging and single-base mutation discrimination of mRNA in living cells. The platform simultaneously incorporates “delivery” and “amplification” functionalities by embedding catalytic hairpin assembly (CHA) modules within a DNA tetrahedral nanoscaffold, where each vertex is anchored with a functional hairpin. This spatially organized configuration allows nanoscale amplification to occur within a confined volume, thereby enhancing reaction efficiency while maintaining molecular stability.

Compared with previously reported DNA nanostructure-based probes, such as tetrahedral molecular beacons or DNA walkers^29–31^, our design integrates localized signal amplification and structural rigidity within a single scaffold, providing superior cellular uptake and stability. In this hierarchical system, three tetrahedra cooperatively execute distinct roles—recognition, initiation, and amplification—and their cascaded arrangement further amplifies fluorescence output. Such an architecture achieves carrier-free, transfection-independent RNA detection with high sensitivity and low background. The 3D@CHA nanoplatform thus resolves the long-standing trade-off between detection sensitivity and intracellular adaptability in nucleic acid imaging.

At the molecular level, the 3D@CHA nanoprobe rationally divides the functional domains of hairpins to balance structural rigidity with dynamic flexibility. The H-AN binds target RNA with high affinity for accurate recognition, while H1/H2 drive strand displacement for cyclic amplification and signal gain. This dual-domain molecular architecture maintains structural integrity while enabling rapid signal turnover, forming the mechanistic foundation for precise and responsive RNA detection in living cells.

Notably, prior CHA-based probes often rely on linear configurations and suffer from slow kinetics and signal leakage^32, 33^. By contrast, the tetrahedral confinement in our system accelerates strand displacement reactions and effectively suppresses background hybridization. Building upon this framework, a single-base mismatch site was deliberately introduced into the H1 stem and combined with a FRET-based ratiometric readout. This design allows subtle base-pairing differences to be translated into distinct optical signatures: perfect matches trigger efficient strand displacement and strong CY3-CY5 FRET signals, whereas mismatches disrupt hairpin opening and attenuate energy transfer. Through this modification, the same 3D@CHA scaffold achieves both real-time RNA imaging and single-base-resolution mutation discrimination within a unified structural context.

These structural and mechanistic innovations endow the 3D@CHA nanoplatform with outstanding performance in live-cell imaging and molecular diagnostics. Without relying on transfection reagents, the nanoprobe spontaneously enters cells and enables dynamic visualization of insulin mRNA localization and transport. Under metabolic modulation, the system captures spatial relocalization and intercellular transfer of insulin mRNA, demonstrating exceptional sensitivity, temporal responsiveness, and biocompatibility. Moreover, validation with the *KRAS^G12D^* mutation model confirmed that the FRET-optimized 3D@CHA nanoprobe could accurately distinguish wild-type and mutant transcripts at single-base resolution, highlighting its utility in early molecular diagnosis and mutation screening over conventional hybridization-based fluorescence probes and molecular beacon systems^34,35^.

With its modular and programmable design, the 3D@CHA nanoplatform provides a versatile molecular framework for diverse RNA-related applications. By simply redesigning the hairpin sequences, the system can be rapidly adapted to target various mRNA, miRNA, or lncRNA species for multiplexed imaging. Compared with CRISPR/Cas13- or aptamer-based RNA sensors that rely on protein components^36–38^, this all-DNA, protein-free system offers higher programmability and avoids cellular immune activation. When combined with protein labeling, metabolic probes, or spatial transcriptomics workflows, the platform could visualize RNA-protein interactions, RNA trafficking, or cellular metabolic networks *in situ*. Integration with high-throughput imaging or single-cell sequencing could further enable spatially resolved transcript profiling and early disease screening, expanding its potential utility in systems biology and precision oncology.

Despite these advances, several limitations remain. Signal-to-noise ratios may still be affected by cellular autofluorescence, and heterogeneous endocytic uptake could lead to partial signal loss in acidic compartments. Moreover, the rapid diffusion and heterogeneous distribution of mRNA continue to pose challenges for quantitative, real-time tracking. Future optimization may involve conjugating the tetrahedral scaffold with cell-penetrating peptides or targeting ligands to improve delivery efficiency^39–41^, and combining the system with super-resolution or real-time tracking microscopy to further enhance spatiotemporal resolution. Such advances could extend the 3D@CHA platform toward *in vivo* imaging and clinical molecular diagnostics.

In summary, this study presents a structurally and functionally integrated 3D@CHA nanoplatform that bridges DNA nanotechnology and RNA biology, enabling real-time visualization and single-base mutation detection of RNA in living cells. By situating our work within the broader context of nucleic acid imaging and mutation analysis, this dual-functional system provides a molecular lens for decoding RNA regulation with base-level precision and offers new avenues for transcript-level diagnostics and precision medicine (Table S5 shown the comparion of perfpormances of recent RNA imaging methods with this strategy.).

## Method

### 1. Synthesis and self-assembly of DNA tetrahedral probes

All DNA and RNA oligonucleotides (Table S1-S4) were synthesized and high-performance liquid chromatography (HPLC)-purified by Sangon Biotech (Shanghai, China). Oligonucleotides were dissolved in hybridization buffer (20 mM phosphate buffer, 0.25 M NaCl, pH 7.4). DNA tetrahedra were assembled by mixing four partially complementary single strands (S1, S3, S4, and S5 to form TDNA-1; S2, S3, S4, and S5 to form TDNA-2; S6, S7, S8, and S9 to form TDNA-3) at equal molar concentrations (4 μM each). The mixture was heated to 95 °C for 10 min and then gradually cooled to room temperature over 3 h to allow stepwise annealing. The assembled tetrahedra were stored at 4 °C until use.

Successful formation of DNA tetrahedra was confirmed by 3.5% native polyacrylamide gel electrophoresis (PAGE) in 1× TBE buffer (pH 8.3) at 80 V for 40 min. Gels were stained with GelRed nucleic acid dye (Biosharp, China) and imaged using a GelDoc XR+ system (Bio-Rad, USA).

### 2. Verification of tetrahedral assembly and CHA activity

To confirm successful assembly, individual strands (S1-S5) were sequentially mixed, annealed, and analyzed by 3.5% native PAGE. Fluorescence measurements were performed using a Synergy HTX multi-mode reader (BioTek, USA) with excitation at 520 nm and emission scanning from 550 to 750 nm.

The fluorescence intensity of each combination was compared to verify stepwise formation of tetrahedral constructs. Three configurations were tested to evaluate CHA efficiency: (1) no CHA reaction (H1+AM only), (2) single-fluorophore CHA (H1 labeled with one F-Q pair), and (3) dual-fluorophore CHA (both H1 and H2 labeled with F-Q pairs). Each system was incubated with AM strand for 3 h at 37 °C, and the peak fluorescence intensity was measured.

### 3. Optimization of reaction conditions

To determine the optimal reaction time, fluorescence intensity was recorded at different incubation periods (0-210 min) under 37 °C. The time at which the signal plateaued was defined as the optimal reaction duration.

To evaluate stability, target-containing and blank samples were stored at 4 °C in the dark and measured daily for 14 days. Fluorescence intensity was calculated relative to the initial signal.

To study temperature dependence, reactions were conducted at 4, 20, 28, 37, 45, 55, 65, 75, 86, and 95 °C for 2 h. After incubation, the difference in peak fluorescence intensity (ΔF = F_target_ − F_blank_) was plotted against temperature.

All experiments were performed in triplicate, and results were expressed as mean ± SEM.

### 4. Analytical performance of the 3D@CHA platform

The analytical performance was evaluated using *Ins1* and *Ins2* mRNAs as model targets. Unless otherwise noted, reactions were performed in hybridization buffer (20 mM phosphate, 0.25 M NaCl, pH 7.4) at 37 °C for 2 h using 100 nM probe.

#### 4.1. Concentration-dependent response and limit of detection

Target mRNAs (0-200 nM) were incubated with probes for 2 h. Fluorescence spectra were recorded, and peak intensities were plotted against target concentrations to construct calibration curves. Linear regression was applied to determine the dynamic range, and the limit of detection (LOD) was calculated as 3σ/k, where σ is the standard deviation of blank measurements and k is the slope of the linear curve.

#### 4.2. Specificity testing

To assess sequence specificity, five mismatched RNA sequences (Mis-1 to Mis-5) were synthesized based on *Ins1*/*Ins2* mRNA, each containing three single-base substitutions. Each sequence (200 nM) was incubated with 100 nM *Ins1*-3D@CHA probe at 37 °C for 2 h, and the fluorescence response was compared with that of the perfectly matched *Ins1*/*Ins2* target.

#### 4.3. Detection in serum samples

Male C57BL/6J mice (8-10 weeks old) were obtained from Vital River and maintained under specific pathogen-free conditions (temperature: 22 ± 2 °C; humidity: 55% ± 5%; 12-h light/dark cycle) at Shanghai Jiao Tong University, with ad libitum access to standard rodent chow and water. After clotting for 30 min, samples were centrifuged at 3,000 rpm for 10 min, filtered (0.22 µm), and stored at −80 °C. Before use, serum was diluted tenfold with hybridization buffer. Target mRNAs (0, 7.5, and 15 nM) were spiked into diluted serum, incubated for 2 h at 37 °C, and fluorescence intensity was recorded. All animal experiments complied with institutional ethical guidelines.

#### 4.4. Dual-domain cooperation

To study synergistic recognition between the anchoring (AN) and amplification (AM) domains, three configurations were tested: AN-only, AM-only, and AN+AM. Each setup was incubated with probe (100 nM) for 2 h at 37 °C, and the resulting fluorescence intensities were compared.

### 5. NUPACK simulation and structural prediction

Secondary structure predictions and free-energy analyses were performed using NUPACK (https://www.nupack.org/). Simulations were conducted at 37 °C with 1 μM strand concentration and 1 M Na⁺ ion concentration. Minimum-free-energy (MFE) structures and base-pairing probabilities were generated for wild-type *KRAS* and mutant *KRAS^G12D^* when the mismatch was positioned at three distinct locations (Positions I, II, III) in the H1 stem. Free-energy differences (ΔΔG) and predicted base-pairing maps were analyzed to evaluate structural stability and mismatch sensitivity.

### 6. Native PAGE validation of simulated complexes

The predicted hybridization outcomes were verified experimentally by 3.5% native PAGE (1× TBE). Reaction mixtures of H1 and *KRAS/KRAS^G12D^*(200 nM each) were incubated at 37 °C for 2 h before electrophoresis. Distinct migration bands confirmed the differing hybridization efficiencies between wild-type and mutant complexes.

### 7. FRET-based probe design and labeling

For mutation-specific detection, a FRET system was established by labeling H1 and H2 strands with CY3 and CY5. All labeled strands were purified by HPLC-purified and verified by MALDI-TOF MS (Sangon Biotech). Fluorescence spectra were collected with excitation at 530 nm and emission scanning from 545 to 750 nm.

This FRET configuration enabled energy transfer-based discrimination between wild-type and mutant *KRAS* targets.

### 8. Cell culture and generation of stable overexpression cell lines

MIN6 cells (mouse insulinoma cells) were purchased from the CAMS Cell Culture Center (Beijing, China). Cells were cultured in Dulbecco’s Modified Eagle Medium (DMEM, Gibco) supplemented with 25.0 mM glucose, 15% fetal bovine serum (FBS), 100 IU/mL penicillin, 100 µg/mL streptomycin, 10.2 mM L-glutamine, and 2.5 mM β-mercaptoethanol. Cultures were maintained at 37 °C in a humidified incubator with 5% CO₂.

293T cells were obtained from the American Type Culture Collection (ATCC) and maintained in DMEM containing 10% FBS and 1% penicillin–streptomycin under the same conditions. For the generation of stable overexpression cell lines, lentiviral particles encoding *KRAS* (wild-type or G12D mutant) or *Ins1*/*Ins2* constructs were used to infect 293T cells at a multiplicity of infection (MOI = 1) in the presence of 6 µg/mL polybrene. After 24 h, the infection medium was replaced with fresh complete DMEM. Forty-eight hours later, cells were selected with 2 µg/mL puromycin for 5–7 days until non-infected controls were eliminated. Stably transduced cells were maintained in 0.5 µg/mL puromycin for subsequent experiments.

### 9. Cytotoxicity assay

Cell viability after DNA tetrahedron exposure was assessed using a CCK-8 kit (Beyotime, Shanghai, China). MIN6 cells were seeded in 96-well plates (1×10⁴ cells/well) and cultured overnight. For concentration testing, cells were treated with 0, 10, 25, 50, 75, and 100 nM DNA tetrahedra for 9 h. For time-dependent testing, cells were incubated with 1 μM tetrahedra for 0, 3, 6, and 9 h. After treatment, 10 μL CCK-8 solution was added and incubated for 1 h at 37 °C. Absorbance at 450 nm was measured with a Synergy HTX reader, and cell viability (%) was calculated as (A_sample_ − A_blank_)/(A_control_ − A_blank_) × 100. All data are presented as mean ± SEM from three independent experiments.

### 10. Confocal fluorescence imaging

293T and MIN6 cells (2×10⁵ cells per dish) were seeded in glass-bottom confocal dishes and incubated for 12 h. The 3D@CHA probe (100 nM) was added and incubated for 8 h at 37 °C. Cells were then washed three times with PBS, stained with Hoechst 33342 (5 µg/mL, 10 min), and imaged using a Leica TCS SP8 confocal microscope (Leica Microsystems, Germany). Intracellular uptake and fluorescence distribution were analyzed at 3, 6, 9, and 12 h using LAS X software (Leica).

### 11. High-to-low glucose switch experiment

To visualize mRNA redistribution during metabolic switching, 293T*^KRAS-G12D^*and MIN6 cells were first incubated with 3D@CHA probe (100 nM) in high-glucose DMEM (4.5 g L^-1^ glucose, 10% FBS) for 8 h to allow probe-mRNA hybridization. The medium was then replaced with low-glucose DMEM (1.0 g L^-1^ glucose, 10% FBS) to initiate metabolic switching. Fluorescence signals were monitored on the Leica TCS SP8 microscope under identical acquisition parameters.

### 12. Western blot (WB) analysis

Total protein was extracted using RIPA lysis buffer (ShareBio, China) supplemented with protease inhibitors and quantified by bicinchoninic acid (BCA) Protein Assay Kit (ShareBio, SB-WB013). Proteins were separated by SDS-PAGE and transferred to PVDF membranes. Membranes were blocked with 5% skim milk (1 h, room temperature) and incubated overnight at 4 °C with primary antibodies: anti-Flag (1:5000, Proteintech, 80801-2-RR), anti-Insulin (1:1000, Abcam, ab181547), and anti-β-actin (1:5000, Proteintech, 66009-1-lg). After washing, membranes were incubated with goat anti-mouse (1:10,000, Proteintech, SA00001-1) or goat anti-rabbit (1:10,000, Proteintech, SA00001-2) secondary antibodies for 1 h at room temperature. Immunoreactive bands were detected by chemiluminescence, quantified using ImageJ, and normalized to β-actin.

### 13. Statistical analysis

All experiments were performed at least three times independently. Data are expressed as mean ± SEM. Statistical analyses were performed using OriginPro 2017.

## Acknowledgements

This work was supported by grants from the National Natural Science Foundation of China (82402436, 92168111, 82230087, 82350123, 82203228), China postdoctoral Science Foundation (2023TQ0220), Postdoctoral Fellowship Program (Grade C) (GZC20231627), Innovative research team of high-level local universities in Shanghai (SHSMU-ZDCX20210802), Shanghai Pilot Program for Basic Research - Shanghai Jiao Tong University (21TQ1400225), 111 project (NO. B21024), Shenyang Science and Technology Plan in 2022 (22-101-0-22) and Key Areas Research and Development Programs of Guangdong Province (2023B1111050009) .

## Author contributions

Z.-G.Z. and X.-Y.M. conceived the project and designed experiments. Y.L., M.M. and X.-Y.M. performed experiments and analyzed the data. X.-Y.M., J.- M.L., and D.-X.L.wrote the manuscript. J.-J.W. and J.- M.L. assisted with experiments. J.-H.Q., L.Z., J.L., L.-P.H., and Y.L. provided critical insights and revised the manuscript. All authors reviewed and approved the final manuscript.

---

**Figure S1.**
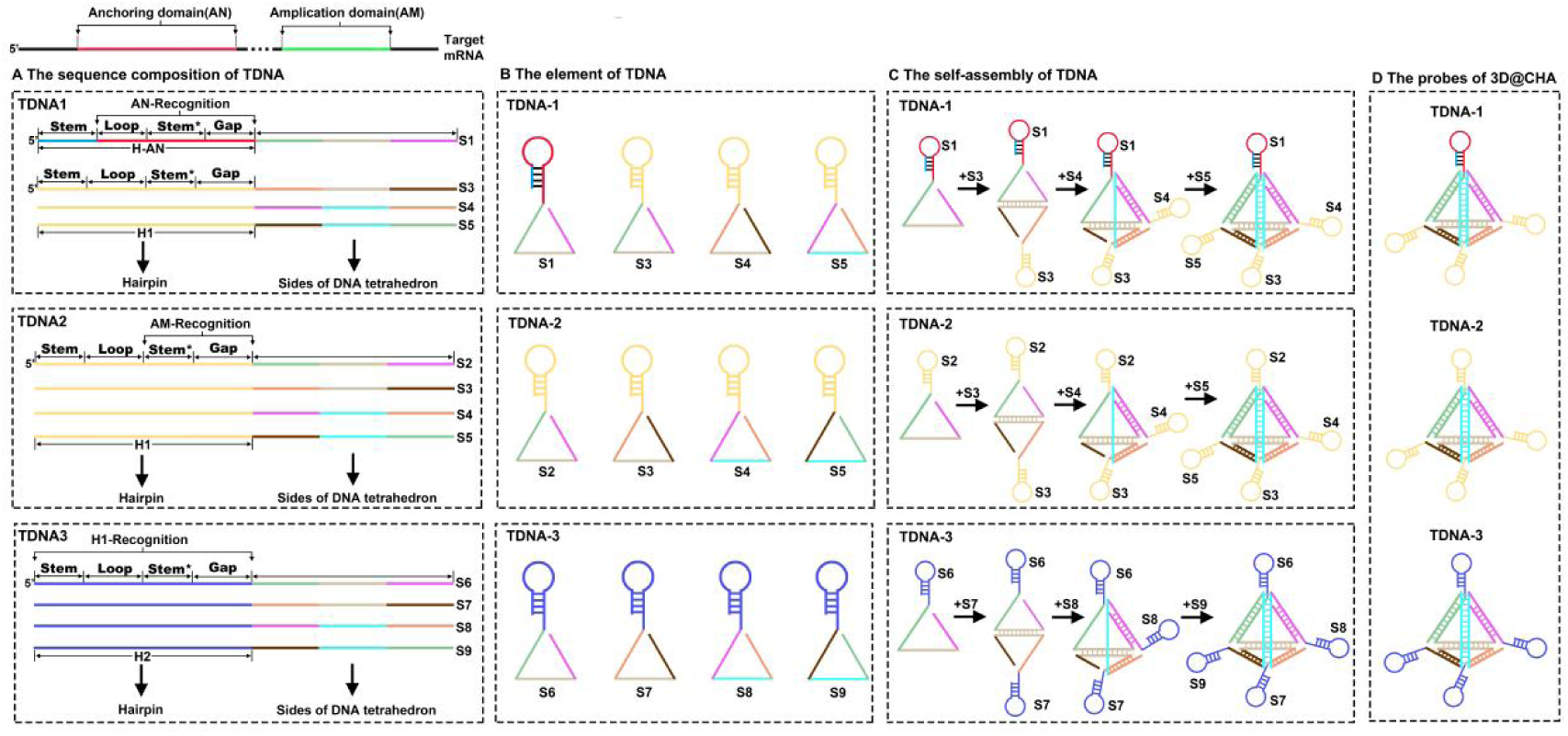
Stepwise assembly and composition of the 3D@CHA probe. (A) The sequence composition of TDNA. Each single-stranded DNA (S1–S9) was designed according to two functional recognition domains of the target mRNA: the anchoring domain (AN) and the amplification domain (AM). The schematic illustrates three tetrahedral DNA units (TDNA-1, TDNA-2, and TDNA-3), each consisting of four single strands that together form both the hairpin structure (for target recognition) and the sides of the DNA tetrahedron (for structural support). TDNA-1: Designed for AN recognition, composed of S1, S3, S4, and S5. The strand S1 carries the anchoring hairpin (H-AN, red) with a Stem–Loop–Stem*–Gap configuration, responsible for specific recognition of the anchoring domain in the target mRNA. TDNA-2: Designed for AM recognition, composed of S2, S3, S4, and S5. The strand S2 forms the amplification hairpin (H1, orange), acting as the initiator for catalytic hairpin assembly (CHA). TDNA-3: Designed for H1 recognition, composed of S6, S7, S8, and S9. The strand S6 carries the complementary amplification hairpin (H2, blue), which reacts with H1 to propagate signal amplification. Each strand within a TDNA unit therefore contributes a portion of the Stem–Loop–Stem*–Gap motif that participates in hairpin formation and simultaneously serves as a structural edge of the DNA tetrahedron, ensuring spatial confinement and geometric stability of the 3D@CHA architecture. In this configuration, “Stem” and “Stem*” represent complementary sequences that hybridize to form the double-stranded stem region of each hairpin (B) Formation of stem-loop-bearing faces of the DNA tetrahedron. Each single strand (S1–S9) folds into a stable Stem–Loop–Gap structure through intramolecular base pairing. The “Stem” and “Stem*” sequences hybridize to form a short double helix, while the loop region provides flexibility and serves as the functional recognition site (H-AN, H1, or H2). In each TDNA, one hairpin-bearing strand (S1 for TDNA-1, S2 for TDNA-2, S6 for TDNA-3) associates with three scaffold strands to create triangular faces representing the sides of a DNA tetrahedron. These triangular faces are shown in distinct colors (green, cyan, and brown) and are functionalized with the corresponding hairpins positioned at one vertex—red for H-AN in TDNA-1, orange for H1 in TDNA-2, and blue for H2 in TDNA-3. (C) Stepwise assembly of individual tetrahedral units. The complete tetrahedral DNA nanostructures are formed through hierarchical hybridization of pre-folded triangular faces, as indicated by the black arrows in the schematic. For each TDNA, strands progressively combine through sticky-end complementarity: In TDNA-1, S1 (bearing the red H-AN) sequentially hybridizes with S3, S4, and S5 to form a fully closed tetrahedral cage containing one H-AN hairpin. In TDNA-2, S2 (bearing the orange H1) joins with S3, S4, and S5, yielding a tetrahedral framework featuring four H1 hairpins that can trigger catalytic hairpin assembly (CHA). In TDNA-3, S6 (bearing the blue H2) hybridizes with S7, S8, and S9 to construct a tetrahedral structure carrying four H2 hairpins, which serve as amplification partners to H1. The sequential “+” steps and directional arrows represent the progressive strand addition process leading to the self-assembly of each tetrahedron, while the loops at the vertices mark the spatial positions of the functional hairpins. (D) Final probe composition and integration into the 3D@CHA architecture. The three functional tetrahedral DNA modules—TDNA-1 (red), TDNA-2 (orange), and TDNA-3 (blue)—are integrated via terminal hybridization to form the complete 3D@CHA probe. In the final nanostructure: TDNA-1 provides the anchoring hairpin (H-AN) for initial target RNA recognition; TDNA-2 provides the H1 hairpins that initiate catalytic amplification; TDNA-3 provides the H2 hairpins that propagate and reinforce signal amplification. Together, these three tetrahedral modules assemble into a three-dimensional catalytic nanoplatform that spatially organizes recognition (H-AN), initiation (H1), and amplification (H2) domains within a confined, rigid framework. This integrated 3D configuration ensures high structural stability, enhanced hybridization kinetics, and efficient fluorescence signal amplification in the 3D@CHA system.

**Figure S2.**
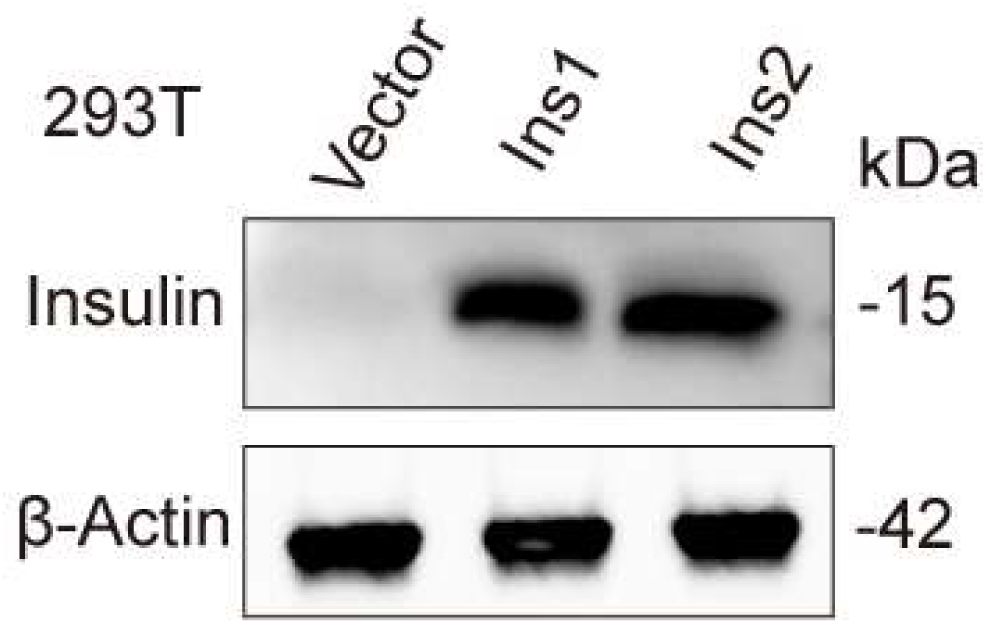
Western blot validation of *Ins*1 and *Ins2* overexpression in 293T cells. 293T cells transfected with *Ins1* or *Ins2* lentiviral constructs exhibited increased insulin expression (∼15 kDa) compared with vector control. β-Actin (∼42 kDa) was used as the loading control. Data are representative of three independent experiments.

**Figure S3.**
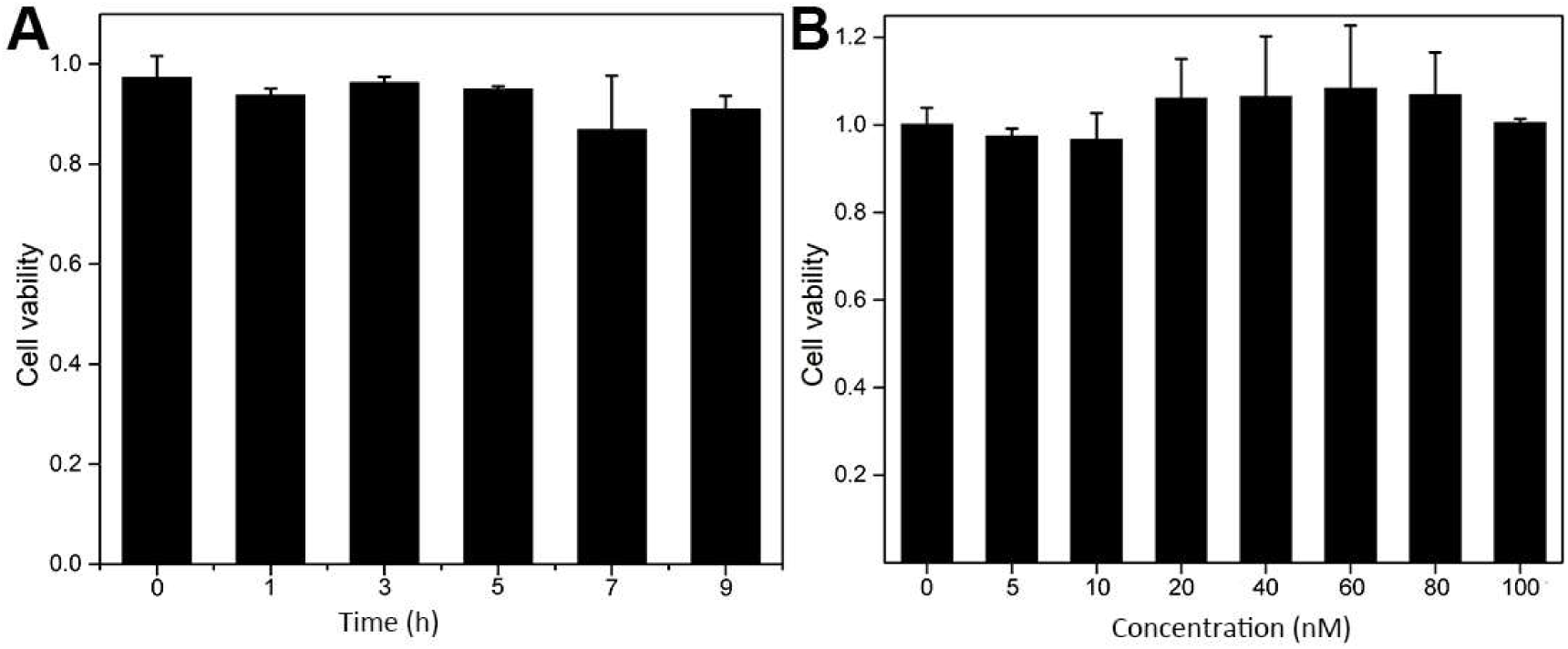
Evaluation of the biocompatibility of DNA tetrahedral probes in 293T cells. (A) Cell viability of 293T cells after incubation with DNA tetrahedra (100 nM) for different time periods (0–9 h). (B) Cell viability of 293T cells after 12 h incubation with DNA tetrahedra at varying concentrations (0–100 nM). Cell viability was assessed using the CCK-8 assay, and absorbance was measured at 450 nm. Data are presented as mean ± SEM from three independent experiments.

**Figure S4.**
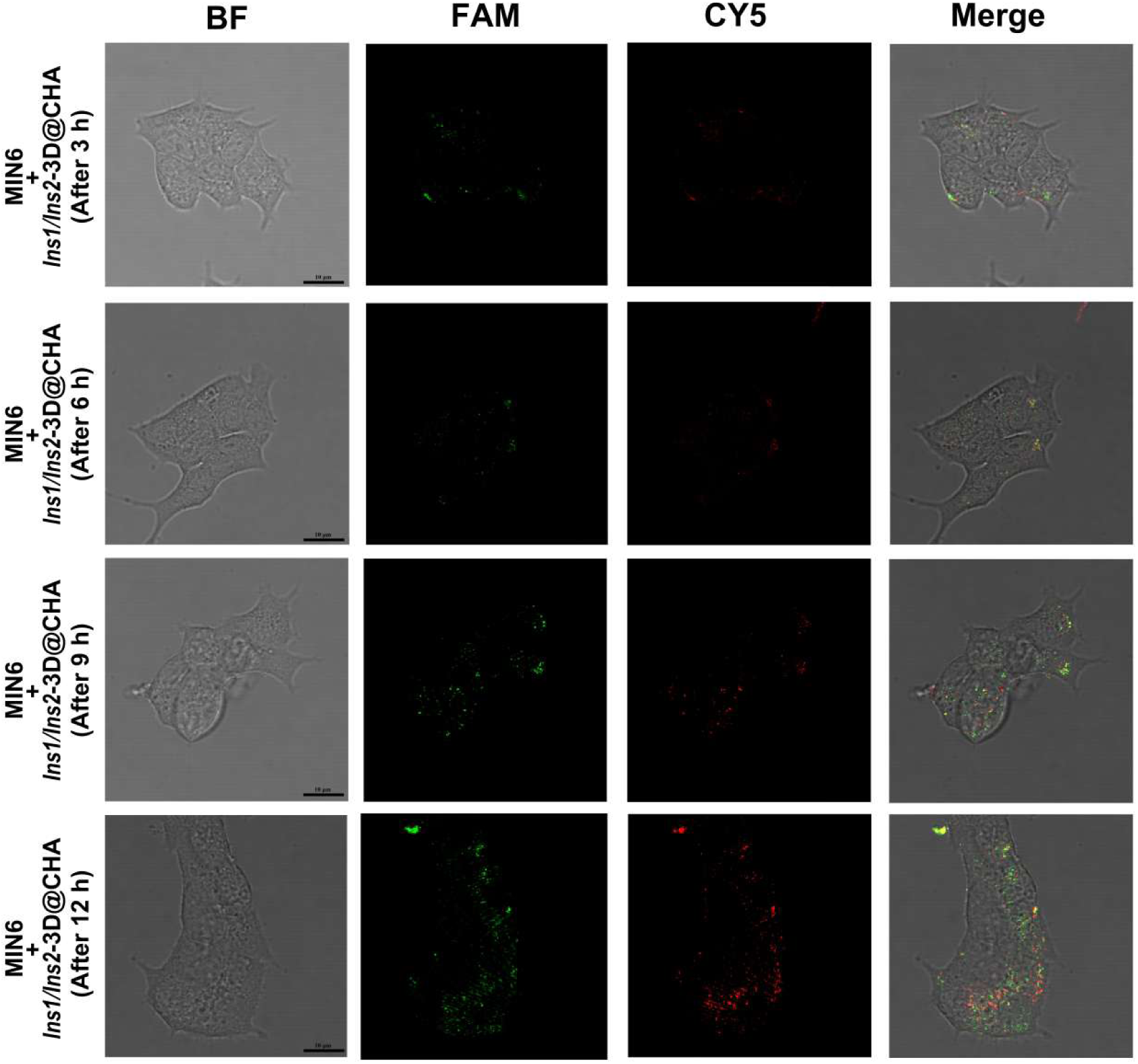
Time-dependent cellular uptake of *Ins1*- and *Ins2*-3D@CHA probes in MIN6 cells. Confocal fluorescence images of MIN6 cells incubated with both *Ins1*-3D@CHA (FAM channel, green) and *Ins2*-3D@CHA (CY5 channel, red) probes for different time periods (3, 6, 9, and 12 h). Bright-field (BF), FAM, and CY5 channels are shown along with merged images. Scale bars, 10 μm.

**Figure S5.**
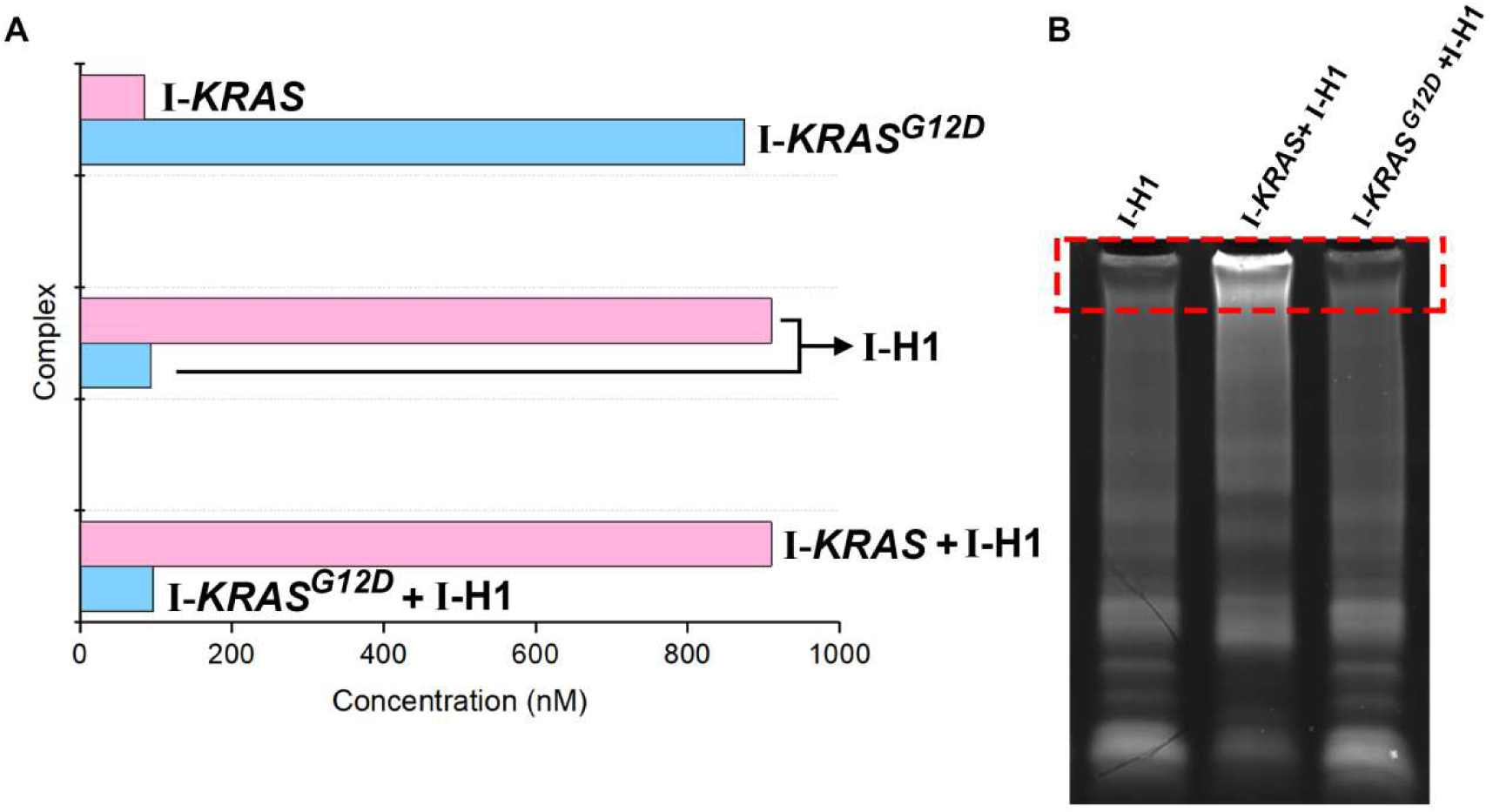
Comparison of reaction efficiency between *KRAS* and *KRAS^G12D^* when the mismatch site is located at Position I. (A) NUPACK-simulated concentrations of various complexes formed between *KRAS/KRAS^G12D^* and H1 when the single-base mismatch site is positioned at Position I of the H1 stem. Perfectly matched *KRAS* readily hybridized with I-H1, generating the *KRAS*-H1 complex, whereas the mismatched *KRAS^G12D^* exhibited reduced binding efficiency and lower complex yield. (B) Native polyacrylamide gel electrophoresis (PAGE) analysis verifying the simulated results. A strong I-H1 band was observed for wild-type *KRAS*, while the corresponding *KRAS^G12D^* + I-H1 lane showed markedly weaker band intensity.

**Figure S6.**
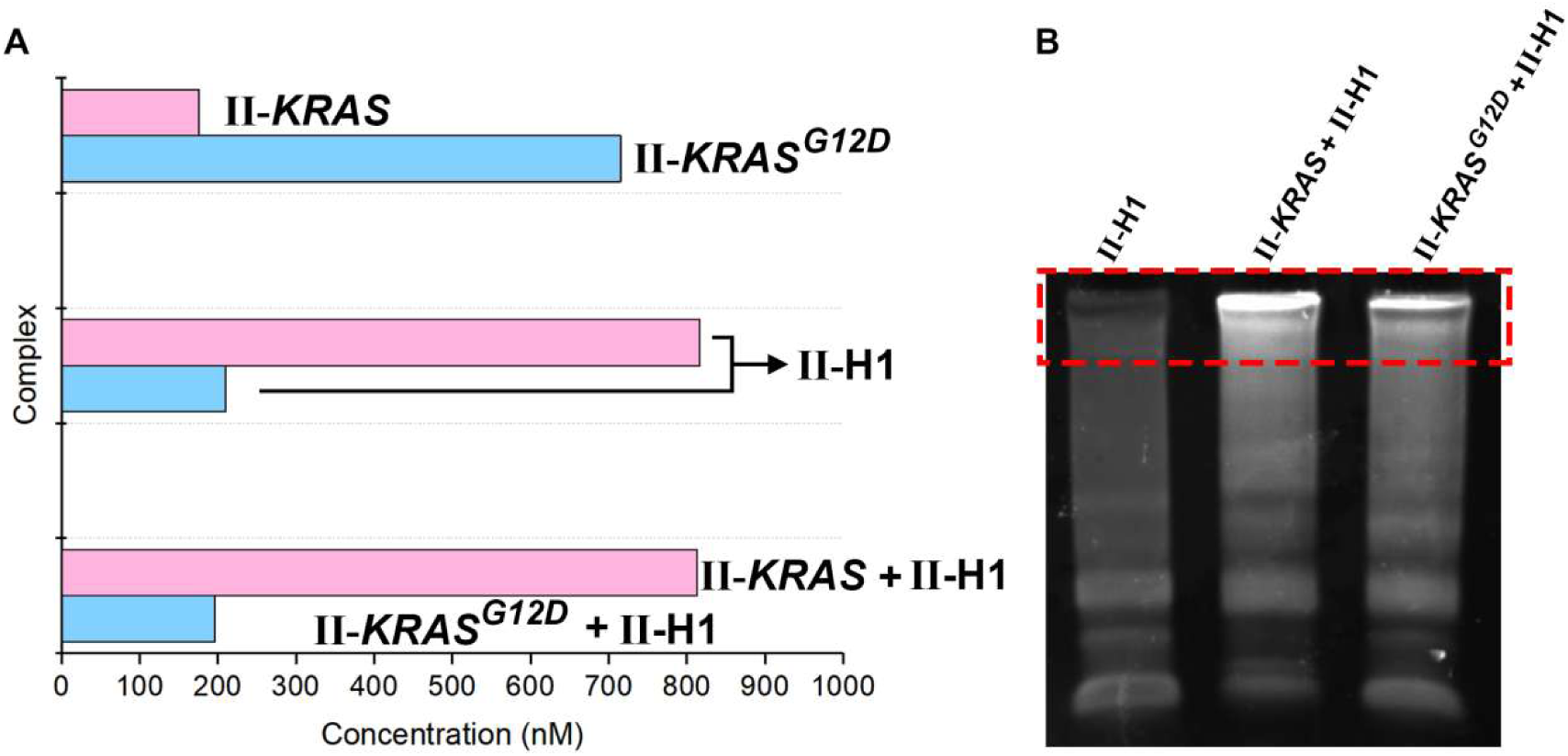
Comparison of reaction efficiency between *KRAS* and *KRAS^G12D^* when the mismatch site is located at Position II. (A) NUPACK-simulated concentrations of reaction intermediates formed between *KRAS/KRAS^G12D^* and H1 when the single-base mismatch site is placed at Position II of the H1 stem. Both *KRAS* and *KRAS^G12D^* could partially hybridize with H1, but the difference in complex formation was smaller than that observed for Position I, suggesting decreased discrimination ability at this site. (B) PAGE analysis confirming the simulation. The II-H1 complex corresponding to wild-type *KRAS* was moderately stronger than that of *KRAS^G12D^*, indicating reduced energetic sensitivity of the Position II region to mismatches.

**Figure S7.**
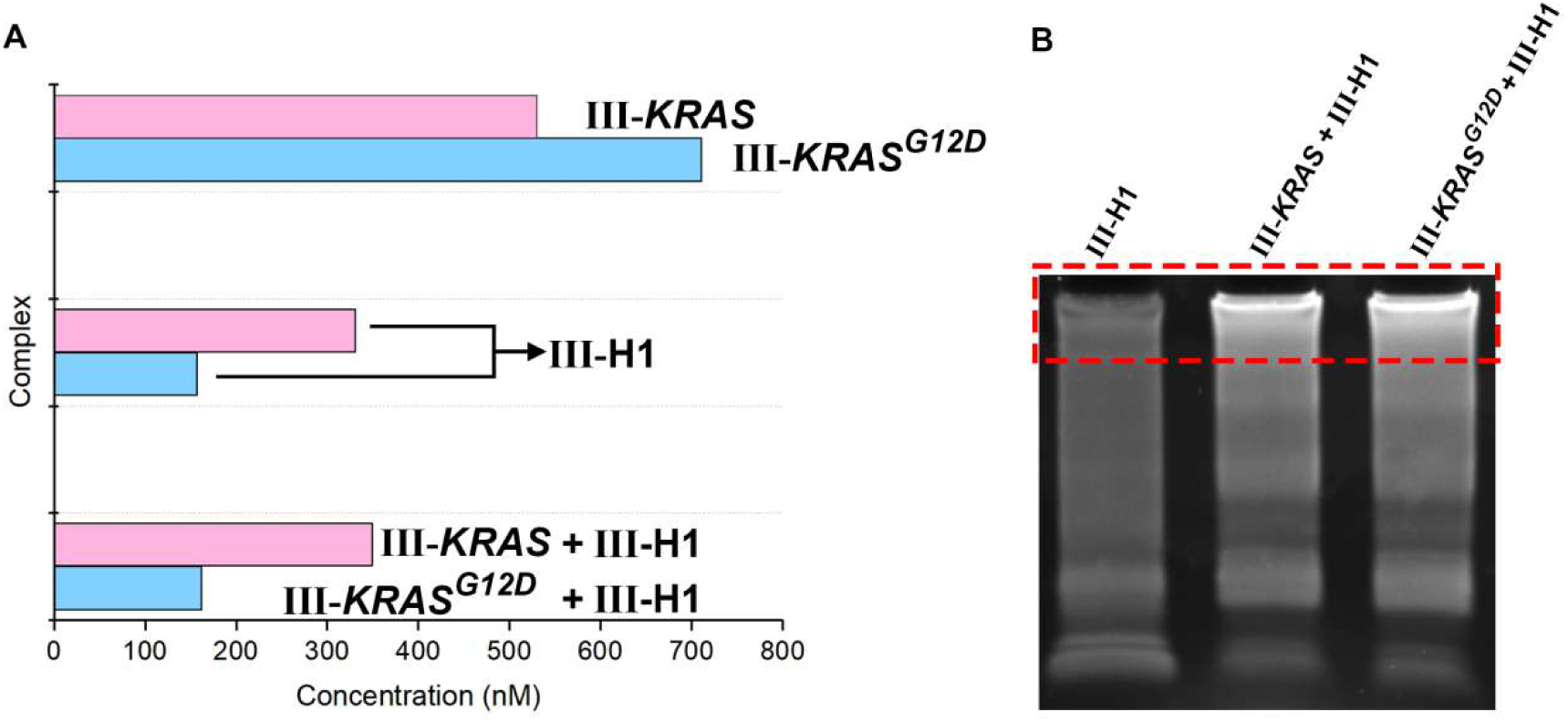
Comparison of reaction efficiency between *KRAS* and *KRAS^G12D^* when the mismatch site is located at Position III. (A) NUPACK-simulated concentrations of hybridization products between *KRAS/KRAS^G12D^* and H1 when the mismatch site is at Position III of the III-H1 stem. The overall complex concentrations were significantly lower than those of Positions I and II, and minimal difference was observed between *KRAS* and *KRAS^G12D^*, indicating that mismatches at the terminal region have limited impact on strand displacement efficiency. (B) PAGE validation of the simulated predictions. Both *KRAS* and *KRAS^G12D^* exhibited faint and comparable III-H1 bands, demonstrating that Position III contributes minimally to mutation discrimination.

**Figure S8.**
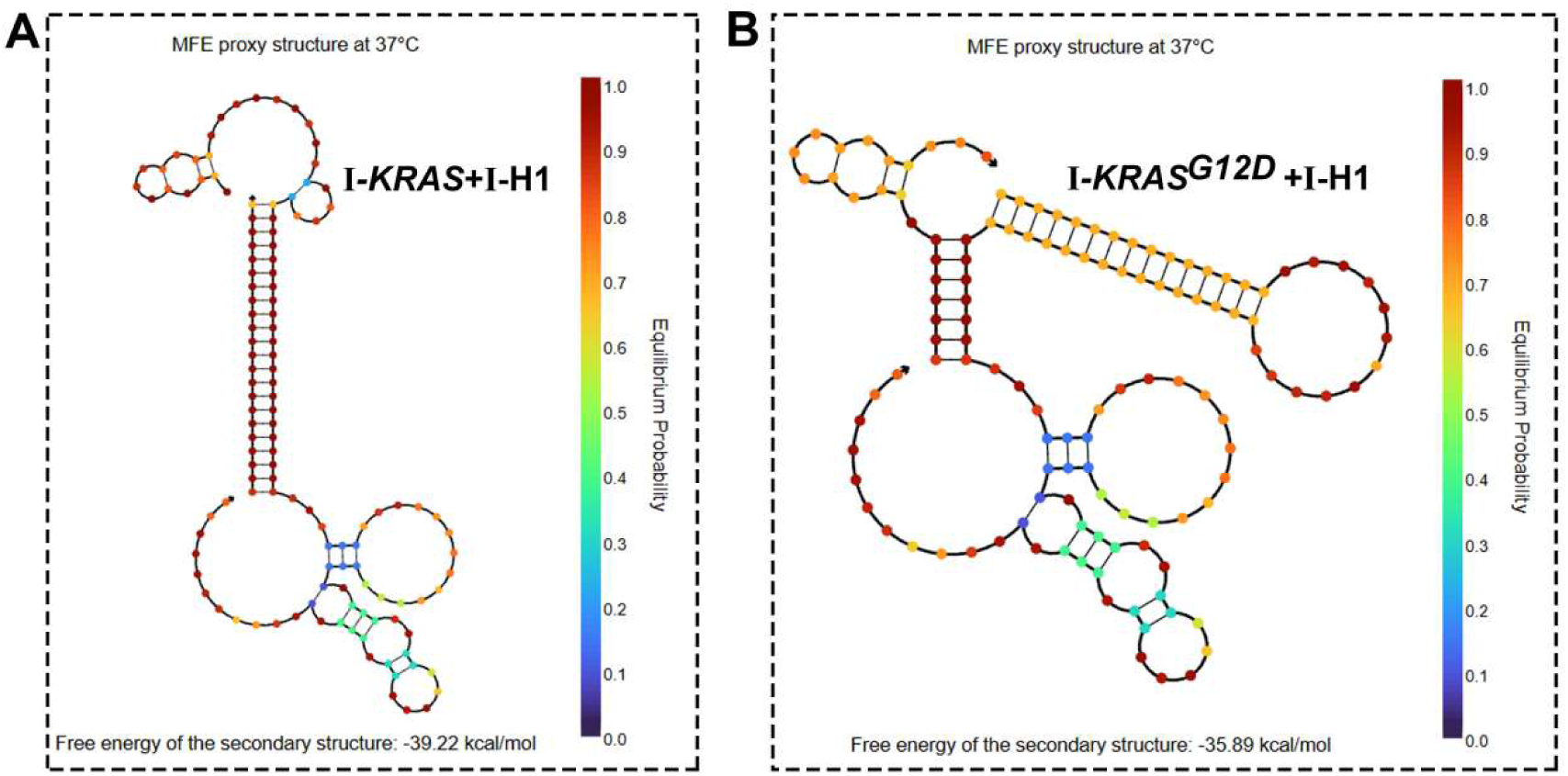
Predicted self-hybridization and secondary structure of *KRAS*-H1 and *KRAS^G12D^*-H1 complexes with the mismatch site at Position I. NUPACK-predicted minimum free energy (MFE) structures and equilibrium base-pairing probabilities of (A) I-*KRAS*-H1 and (B) I-*KRAS^G12D^*-H1 complexes when the single-base mismatch site is located at Position I (the initiation region of the H1 stem). Color coding indicates base-pairing equilibrium probabilities (from 0.0, blue, to 1.0, red). The predicted free energy of the secondary structure is -39.22 kcal/mol for I-*KRAS*-H1 and -35.89 kcal/mol for I-*KRAS^G12D^*-H1, suggesting that the perfectly matched I-*KRAS*-H1 complex adopts a more stable conformation with a lower free energy. In contrast, the I-*KRAS^G12D^*-H1 complex shows partial disruption of the stem region due to the mismatch, leading to a less stable structure and reduced hybridization efficiency.

**Figure S9.**
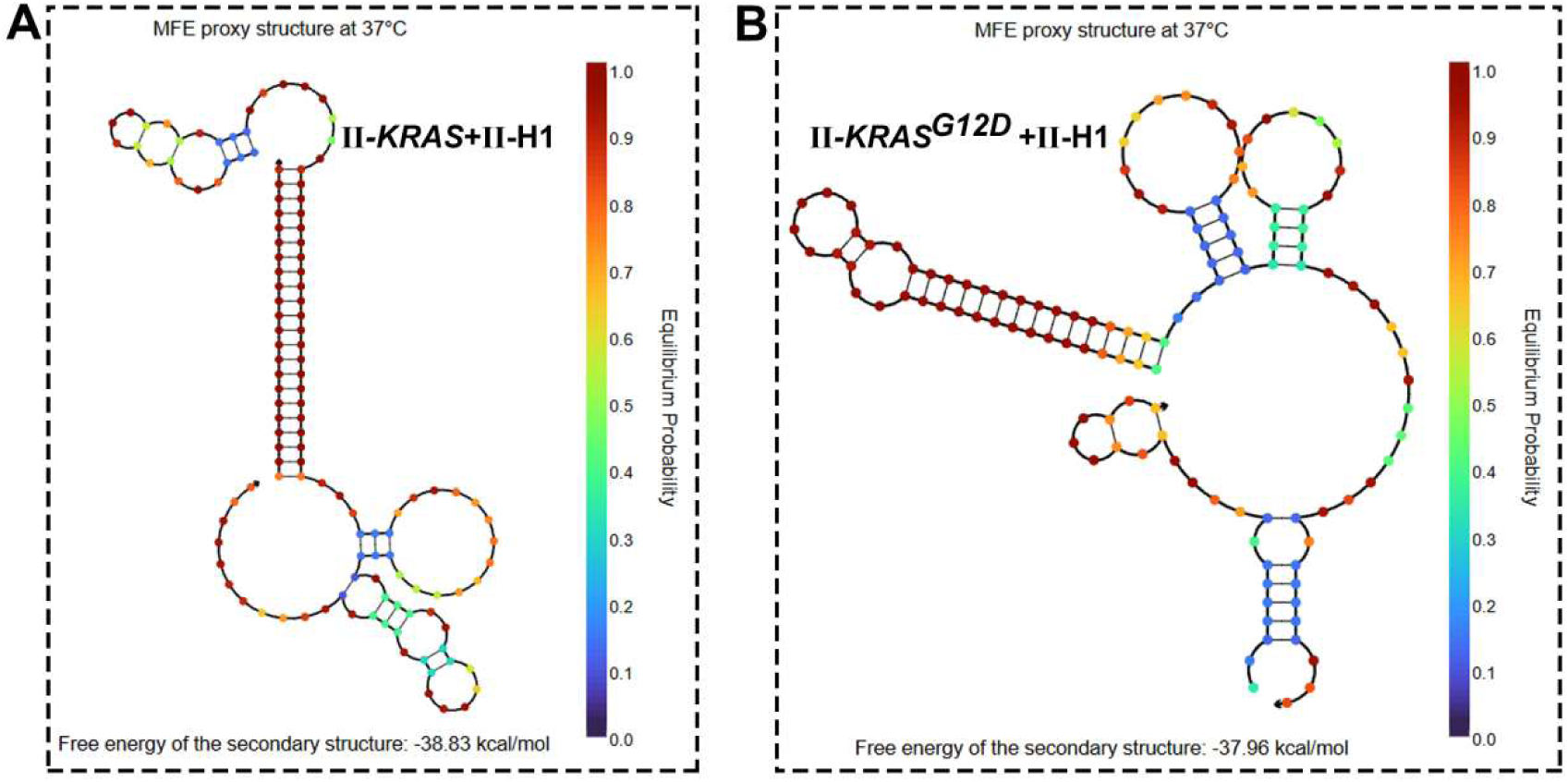
Predicted self-hybridization and secondary structure of *KRAS*-H1 and*KRAS^G12D^*-H1 complexes with the mismatch site at Position II. Secondary Structures of (A) *KRAS* and (B) *KRAS^G12D^* bound to II-H1 when the mismatched site is at Position II. NUPACK-predicted MFE structures and equilibrium base-pairing probabilities of (A) II-*KRAS*-H1 and (B) II-*KRAS^G12D^*-H1 when the mismatch site is located at Position II (the middle region of the H1 stem). Color scale represents the equilibrium probability of each base pair (0.0 = blue, 1.0 = red). The predicted free energy of the secondary structure is -38.83 kcal/mol for II-*KRAS*-H1 and -37.96 kcal/mol for II-*KRAS^G12D^*-H1. The slight difference in free energy indicates that mismatches at Position II induce limited structural perturbation. However, minor local loop distortions are visible in the II-*KRAS^G12D^*-H1 complex, reflecting weaker hybridization at the mismatch site and moderately reduced binding stability compared with the wild-type II-*KRAS*-H1 structure.

**Figure S10.**
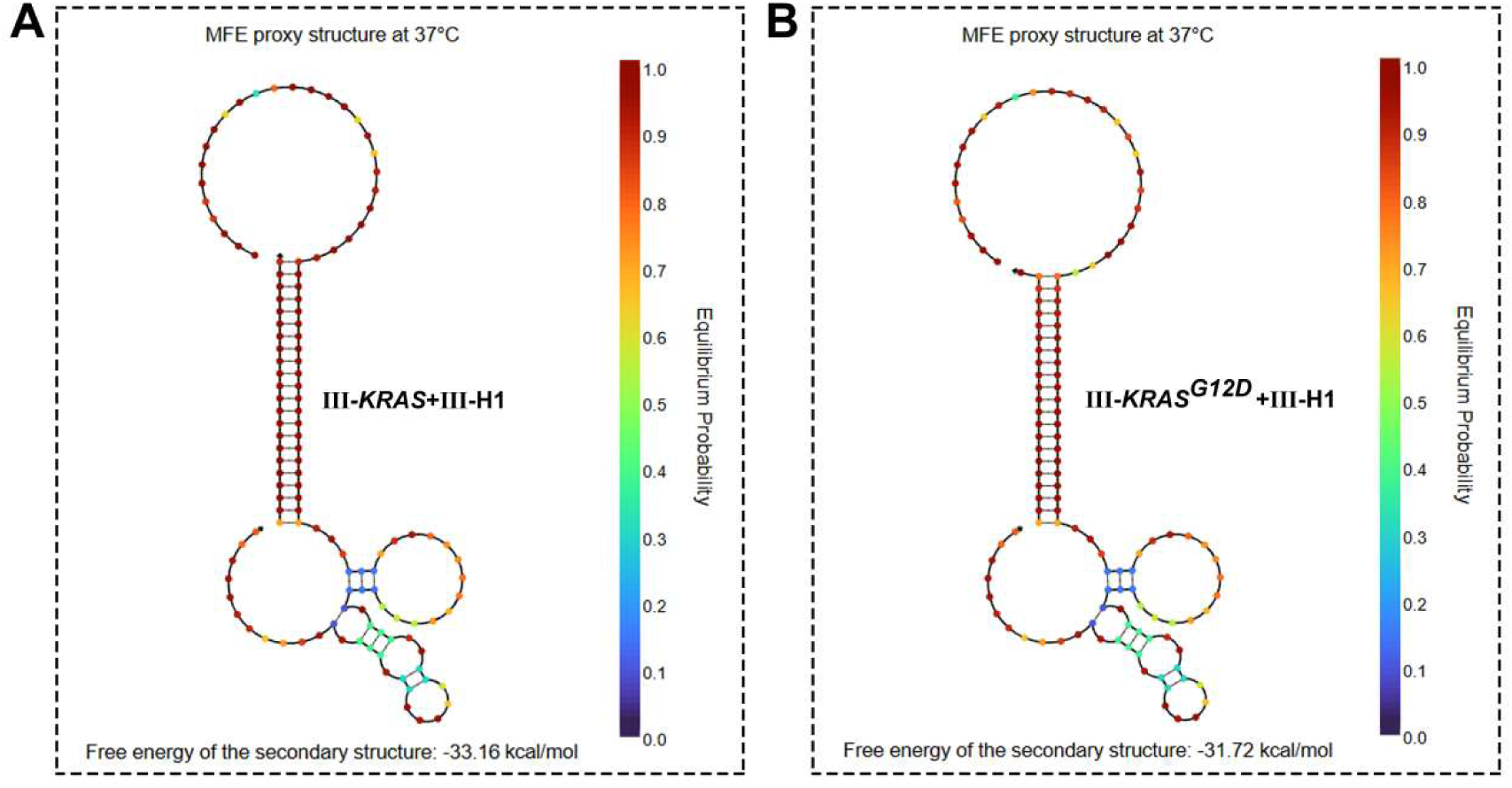
Predicted self-hybridization and secondary structure of *KRAS*-H1 and *KRAS^G12D^*-H1 complexes with the mismatch site at Position III. NUPACK-predicted MFE structures and equilibrium base-pairing probabilities of (A) III-*KRAS*-H1 and (B) III-*KRAS^G12D^*-H1 when the mismatch site is located at Position III (the terminal region of the H1 stem). Color scale represents base-pair equilibrium probabilities (0.0 = blue, 1.0 = red). The predicted free energy values are -33.16 kcal/mol for III-*KRAS*-H1 and -31.72 kcal/mol for III-*KRAS^G12D^*-H1, indicating the weakest structural stability among the three tested positions. Both complexes maintain an overall intact stem-loop configuration, and the free energy difference between wild-type and mutant forms is minimal, suggesting that terminal mismatches exert limited influence on reaction kinetics and mutation discrimination capability.

**Figure S11.**
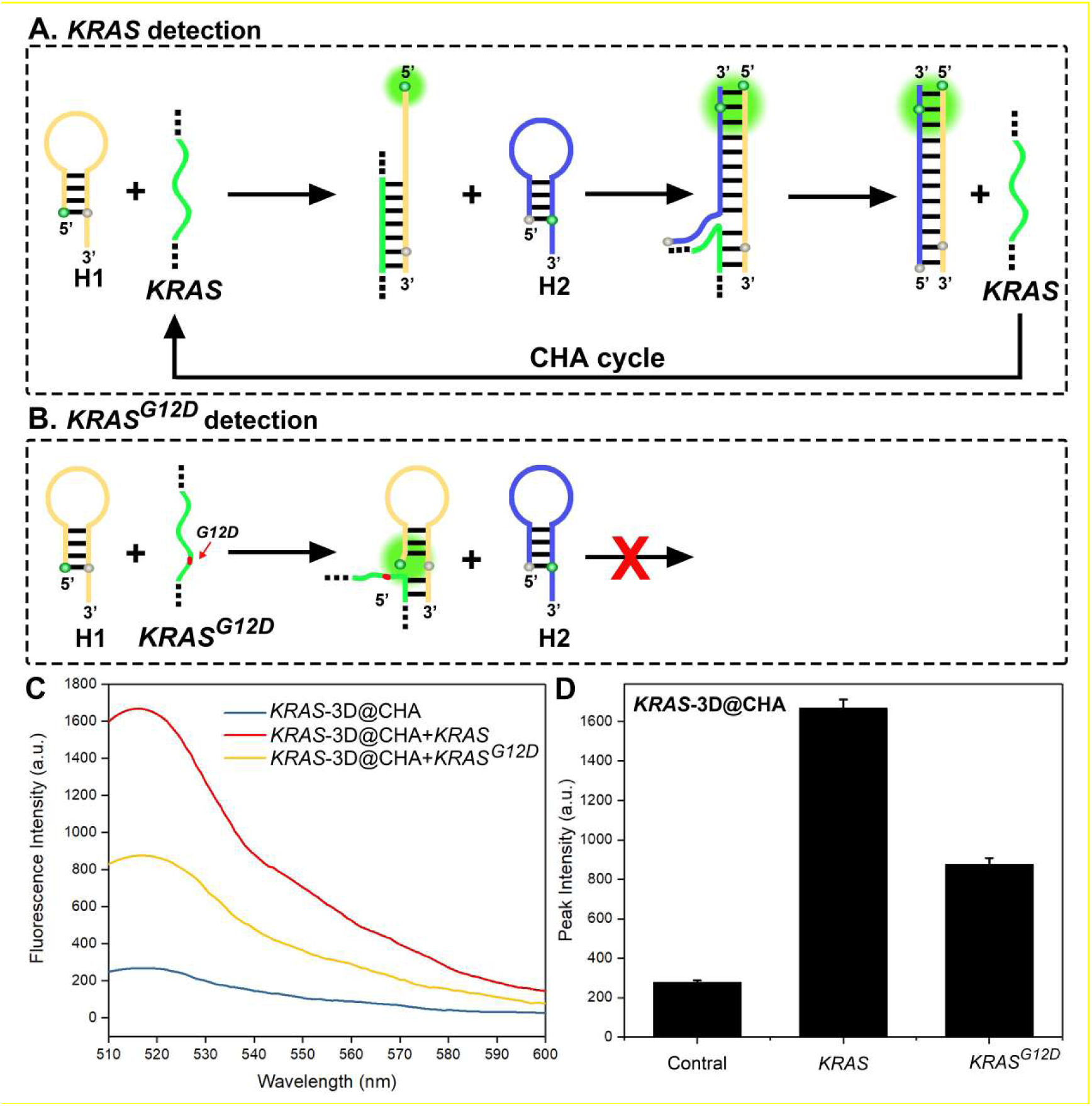
Fluorescence turn-on mechanism and single-base mutation discrimination of the *KRAS*-3D@CHA probe. (A) Schematic illustration of the CHA process triggered by wild-type *KRAS* mRNA. The *KRAS*-3D@CHA probe consists of two hairpin strands, H1 (orange) and H2 (blue), embedded within a three-dimensional tetrahedral DNA framework. Upon hybridization with the target *KRAS* mRNA, the 5′ region of H1 binds to the complementary sequence, inducing hairpin opening and exposing a toehold for strand displacement with H2. This reaction generates an H1–H2 duplex and releases the *KRAS* trigger to initiate subsequent CHA cycles, thereby producing a strong fluorescence signal (“turn-on” effect). wild-type and mutant forms is minimal, suggesting that terminal mismatches exert limited influence on reaction kinetics and mutation discrimination capability. (B) Detection mechanism for the *KRAS^G12D^*mutant mRNA. The single-base substitution (GGT→GAT, marked in red) at the hybridization site introduces a mismatch that destabilizes the H1–*KRAS* interaction. Consequently, the hairpin fails to open properly, and the CHA cycle cannot proceed efficiently. The red cross indicates termination of the amplification process caused by mismatch-induced hybridization failure, resulting in minimal fluorescence output. This mechanism enables precise discrimination between wild-type and mutant *KRAS* transcripts at single-nucleotide resolution. (C) Fluorescence emission spectra of the *KRAS*-3D@CHA system. Spectra were recorded in the absence of target RNA (control, blue), in the presence of wild-type *KRAS* mRNA (red), or in the presence of *KRAS^G12D^* mRNA (yellow). A pronounced fluorescence enhancement was observed only with wild-type *KRAS*, confirming target-dependent activation of the CHA reaction. (D) Quantitative comparison of fluorescence peak intensities. *KRAS* mRNA produced a strong “turn-on” response, whereas the *KRAS^G12D^* mutant exhibited approximately half the fluorescence intensity of the wild-type *KRAS* signal. Error bars represent mean ± SEM from three independent experiments.

**Figure S12.**
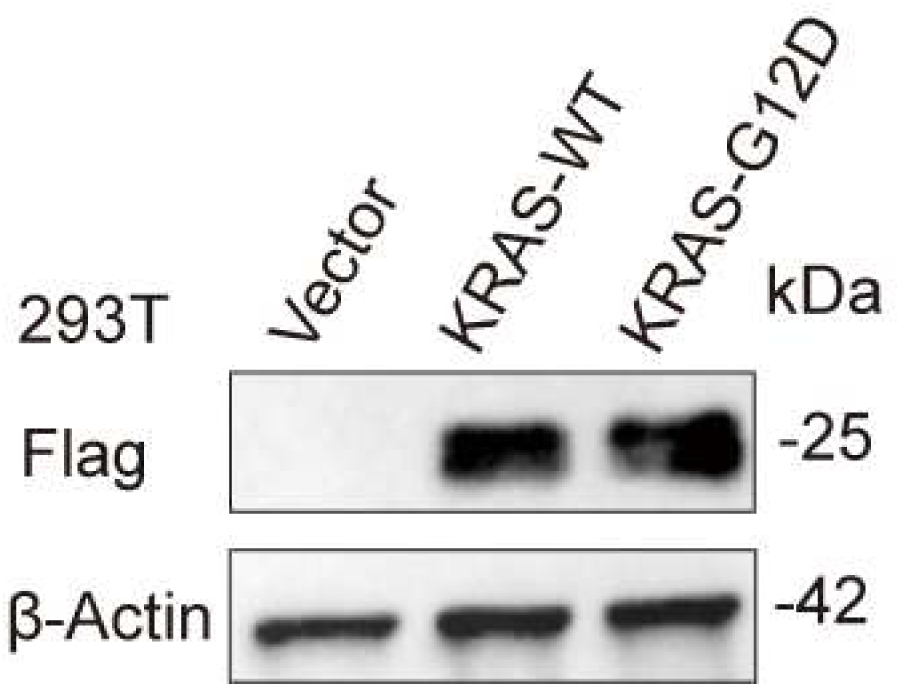
Western blot validation of *KRAS* overexpression in 293T cells. 293T cells were transfected with lentiviral constructs encoding wild-type *KRAS* (*KRAS-WT)* or the G12D mutant (*KRAS-G12D*), with an empty vector as control. Cell lysates were analyzed by Western blot using an anti-Flag antibody to detect *KRAS* expression (∼25 kDa). β-Actin (∼42 kDa) served as the loading control.

**Table S1.**
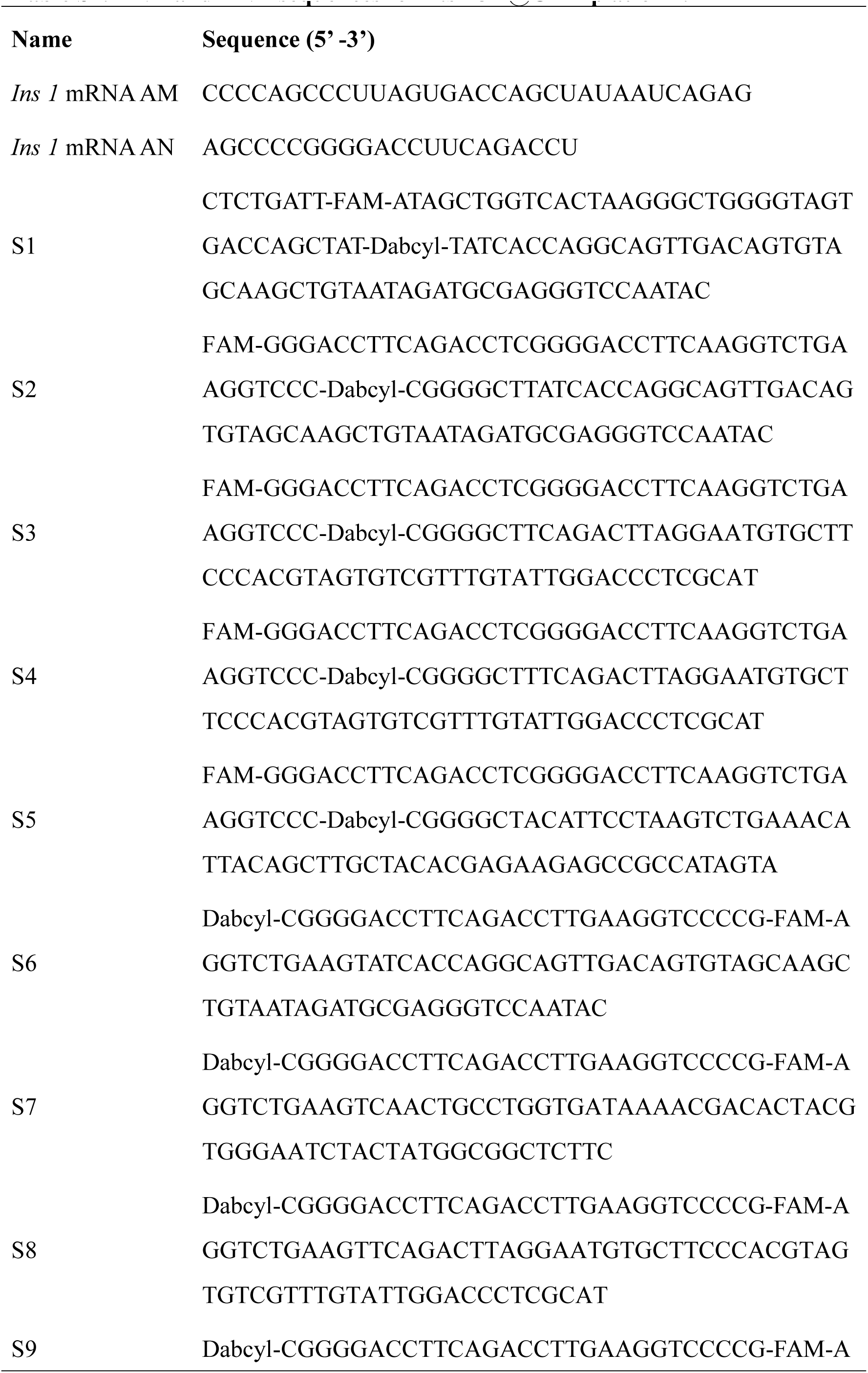
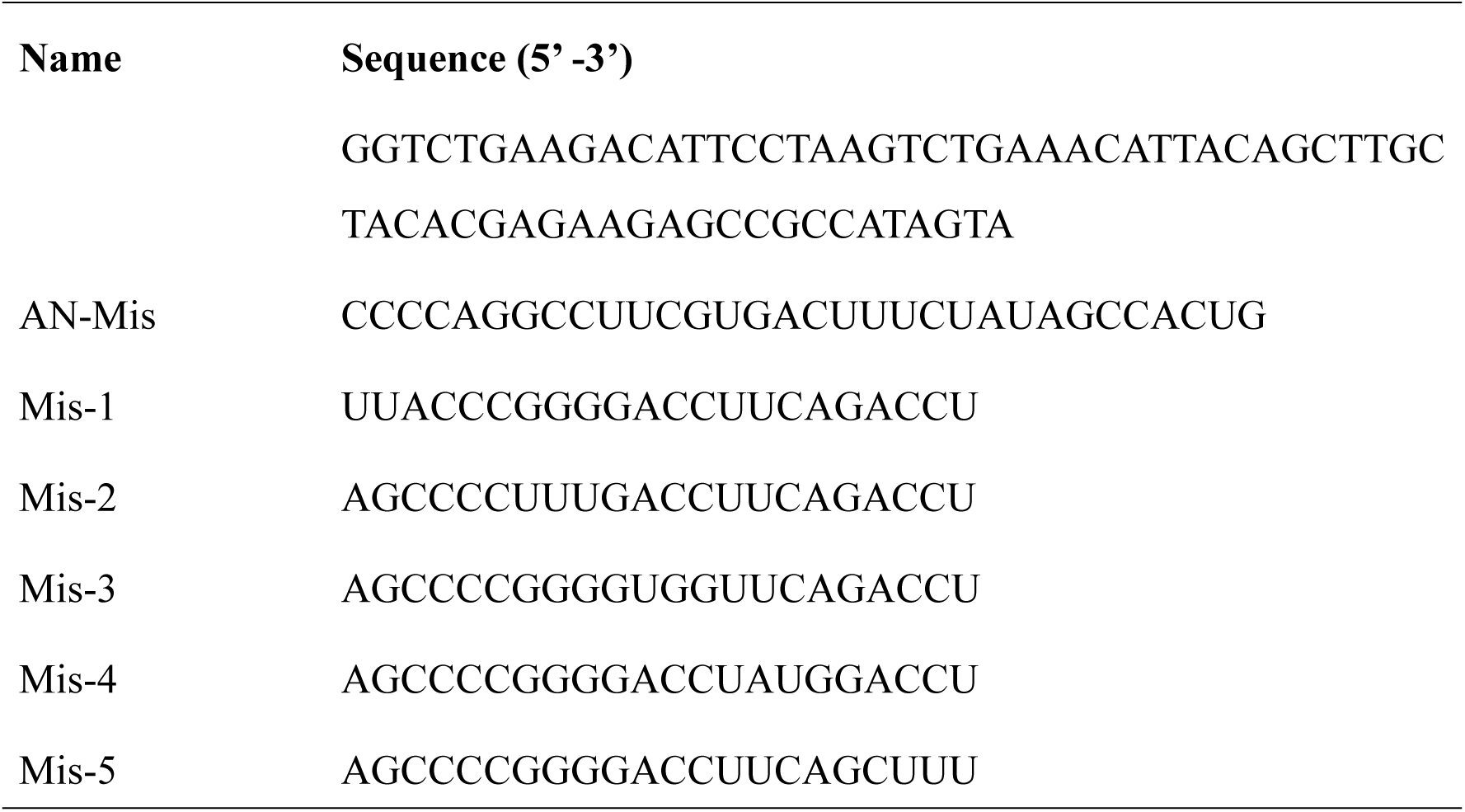
DNA and RNA sequences for *Ins1*-3D@CHA platform.

**Table S2.**
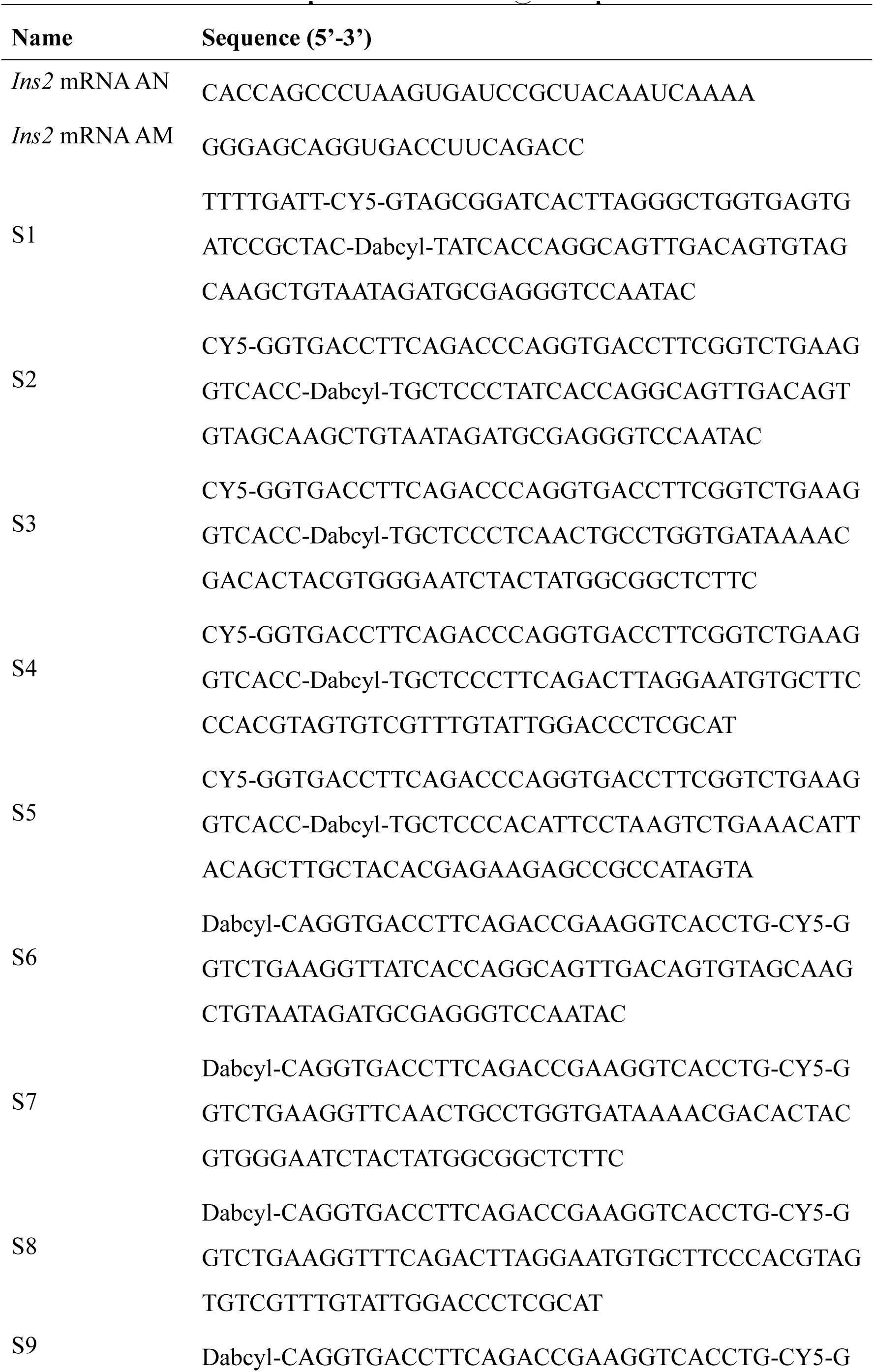
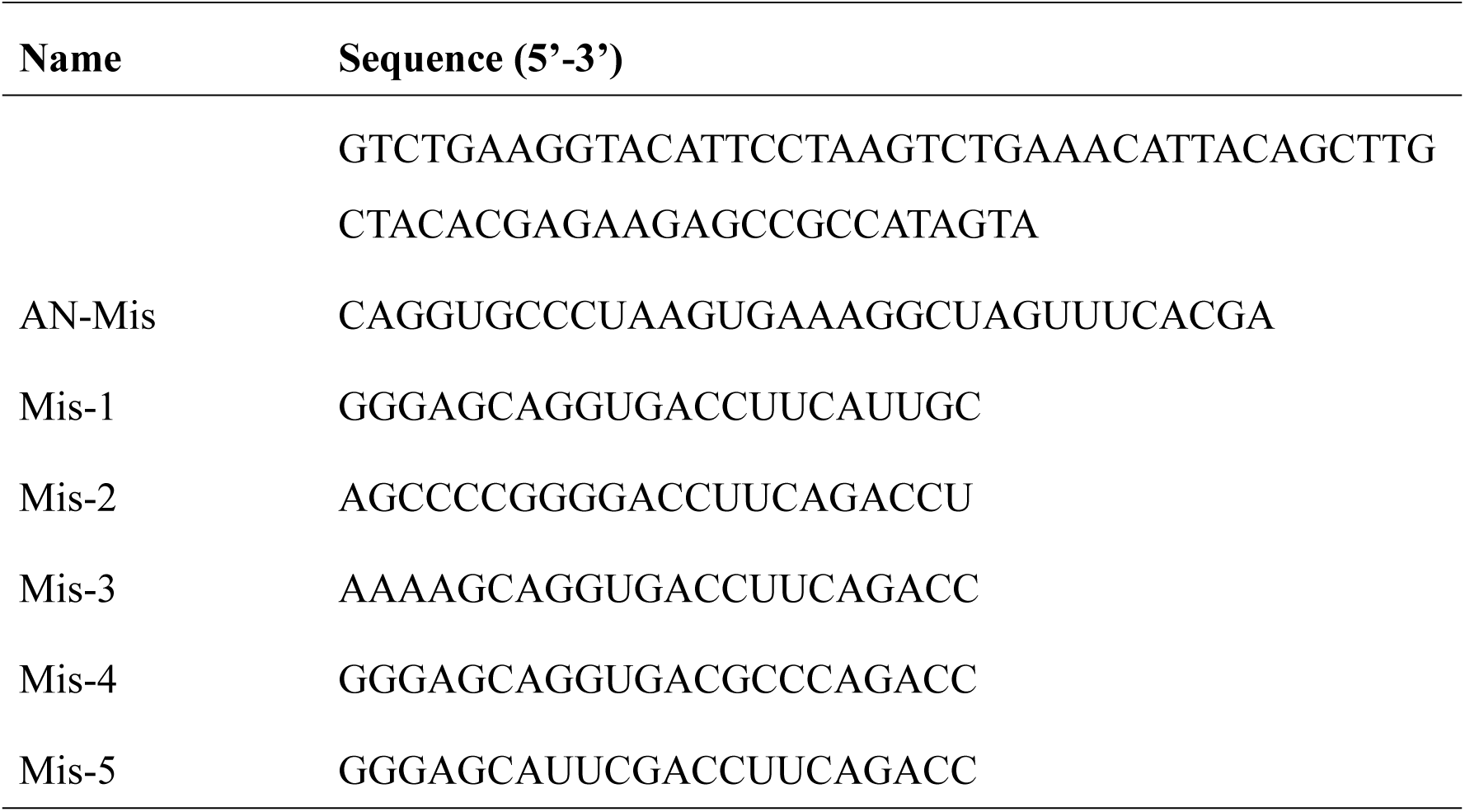
DNA and RNA sequences for *Ins2*-3D@CHA platform.

**Table S3.**
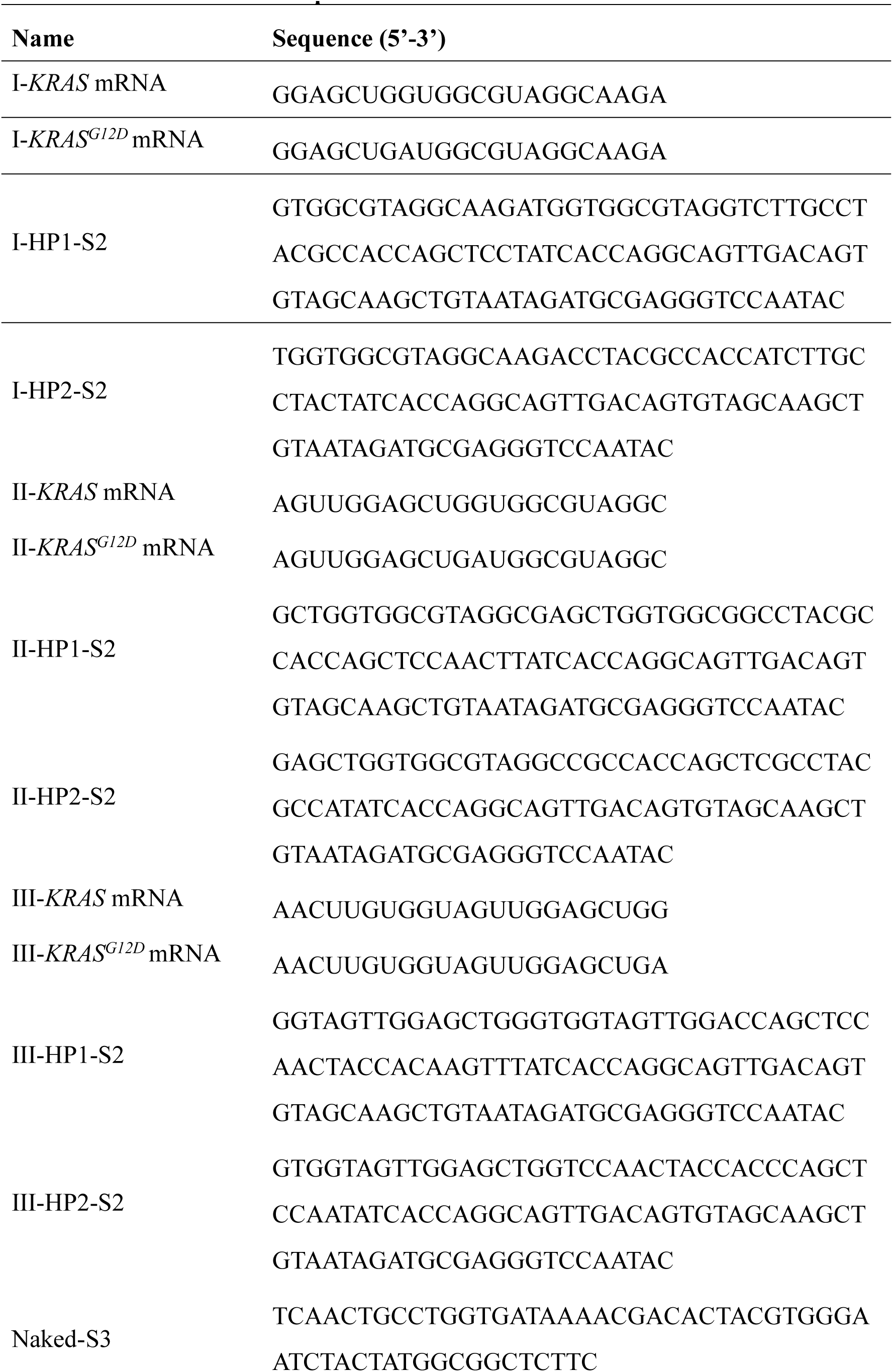
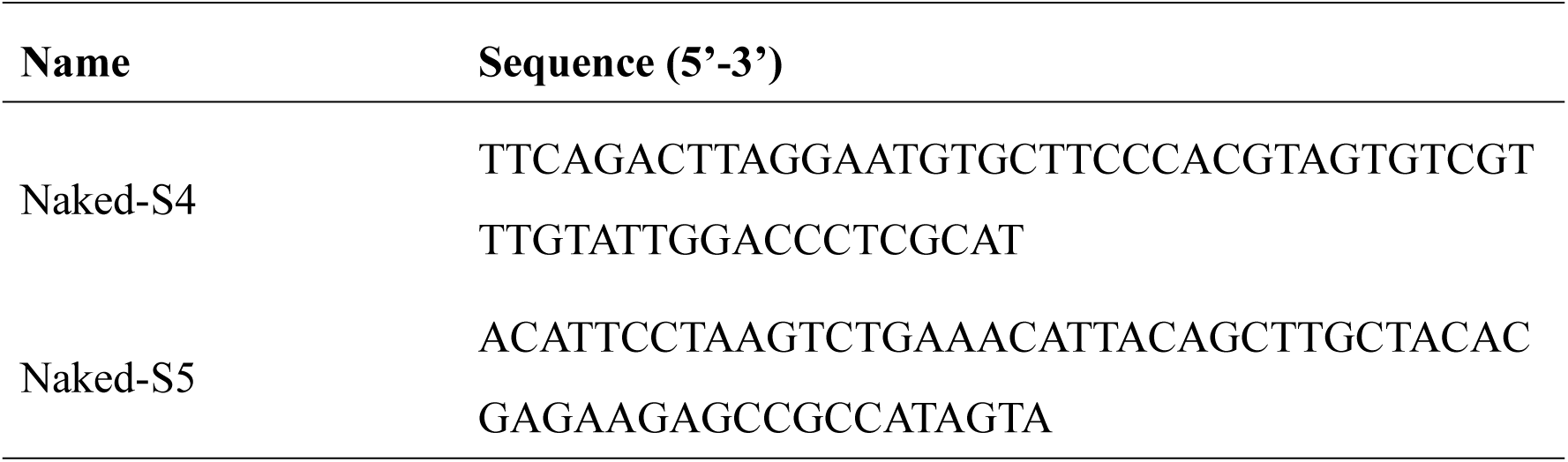
DNA and RNA sequences for the location of mismatch base.

**Table S4.**
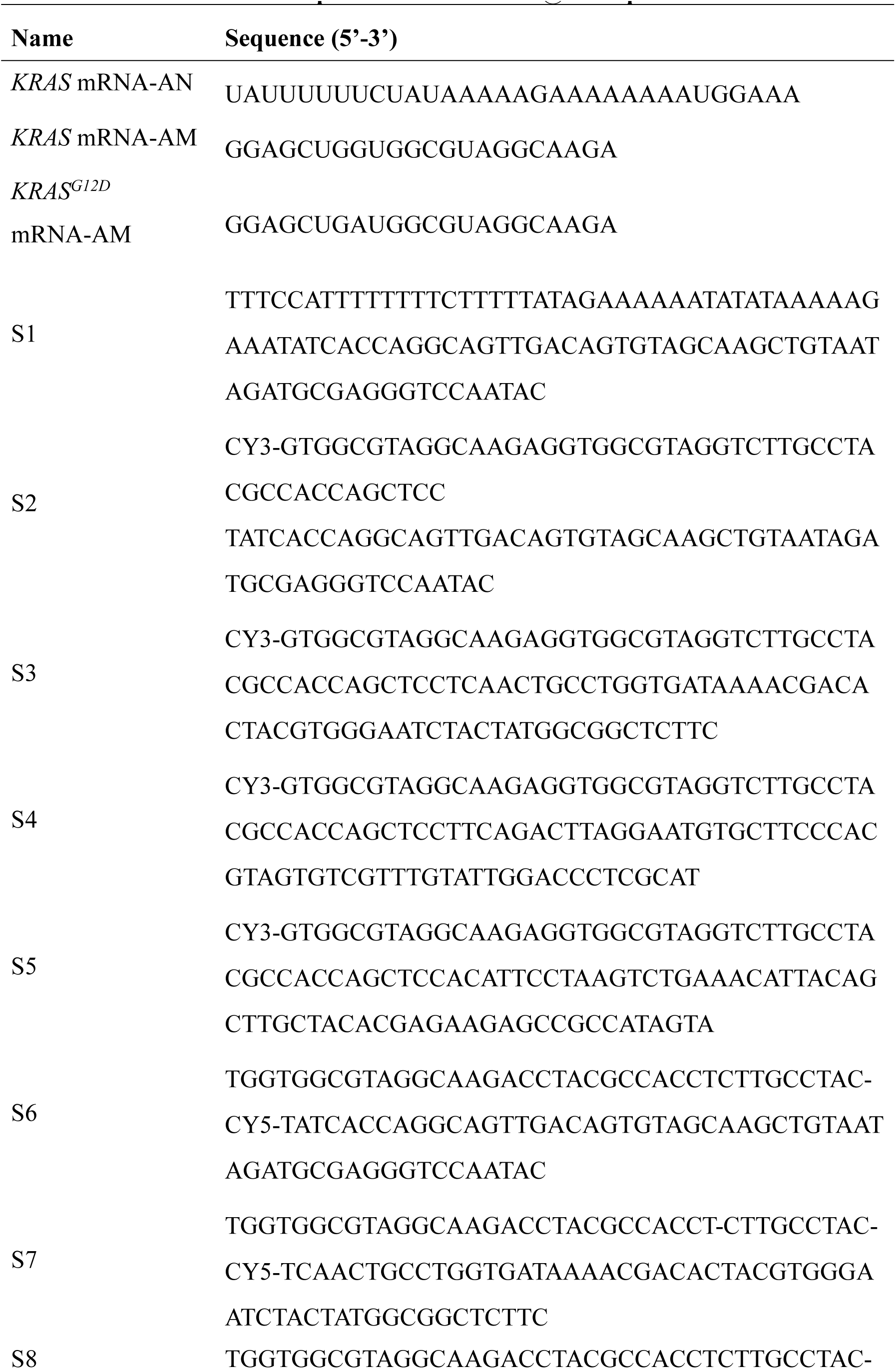
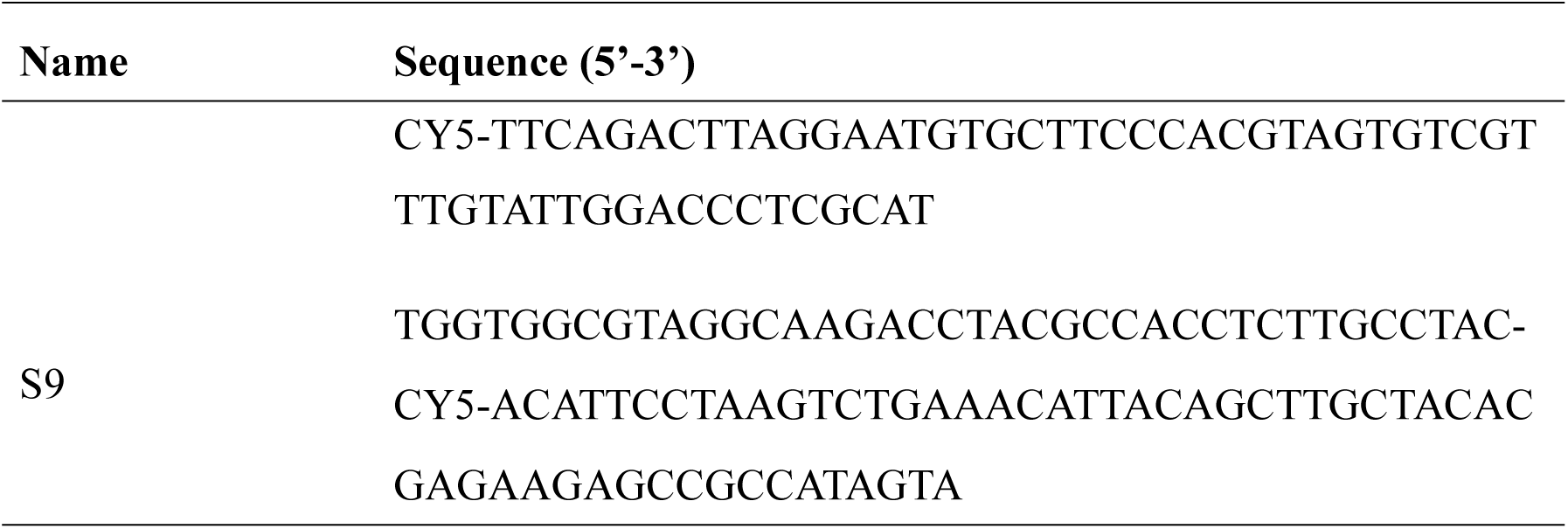
DNA and RNA sequences for *KRAS*-3D@CHA platform.

**Table S5.**
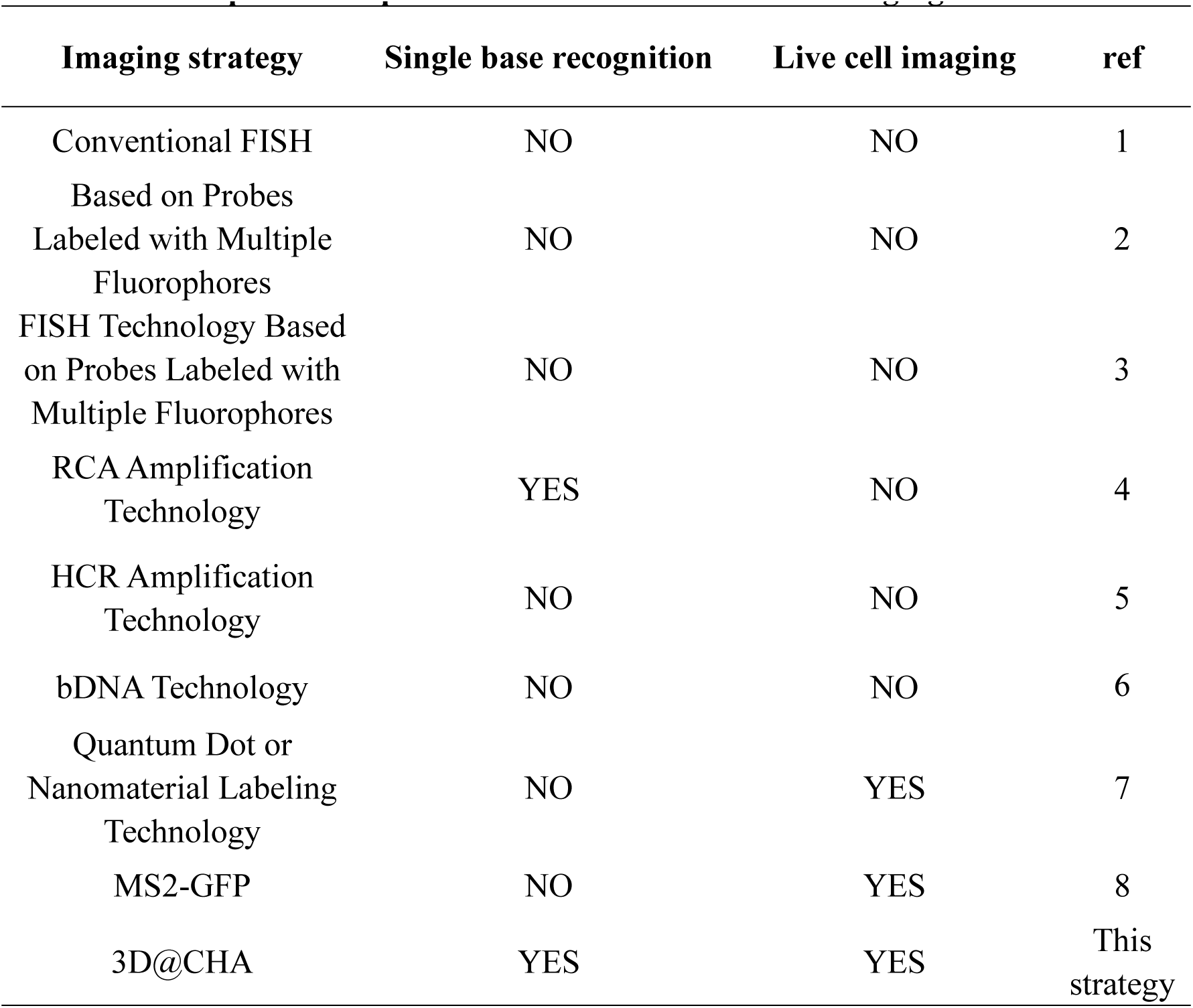
Comparison of performances of recent mRNA imaging methods.

**Table S6.**
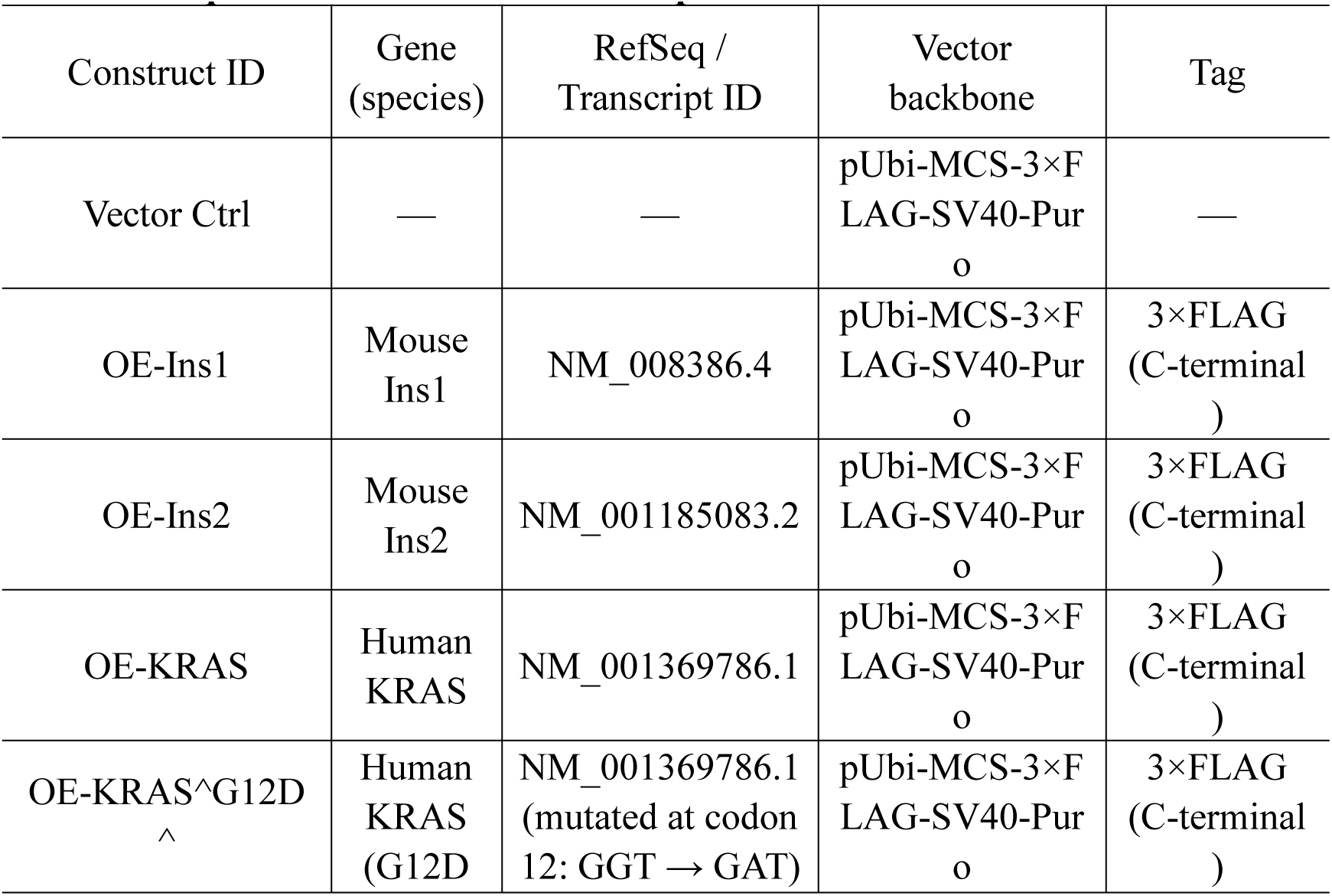
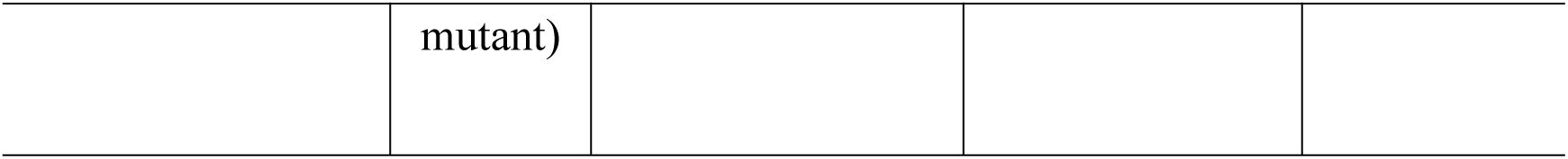
Sequence information of overexpression constructs used in 293T cells.

**Table S7.** The reagents used in this work.

| Reagent | Company |
| --- | --- |
| NaCl | Aladdin Biotech Co., Ltd. (Shanghai, China) |
| NaH<sub>2</sub>PO<sub>4</sub>·2H<sub>2</sub>O | Aladdin Biotech Co., Ltd. (Shanghai, China) |
| Na<sub>2</sub>HPO<sub>4</sub>·12H<sub>2</sub>O | Aladdin Biotech Co., Ltd. (Shanghai, China) |
| 20 bp DNA ladder | Takara Biotechnology (Dalian, China) |
| gel loading buffer | Solarbio Life Sciences (Beijing, China) |

